# DESI-MS-Based Analysis of Drug Distribution in Human Renal Cystic Tissue Using the Chorioallantoic Membrane (CAM) as a 3D In Vivo Model

**DOI:** 10.64898/2026.07.01.735776

**Authors:** Antonia Hehemann, Jan Schueler, Simon Heckscher, Verena Groß, Matthias May, Barbara Nuebel, Bernd Wullich, Bjoern Buchholz, Jens M. Werner, Jonathan Jantsch, Wolfram Gronwald, Zoltan Takats, Peter J. Oefner, Katharina M. Schmidt, Silke Haerteis, Katja Dettmer

**Author notes:** Corresponding authors: **Silke Haerteis** - Institute of Molecular and Cellular Anatomy, University of Regensburg, Regensburg, Germany, **Katja Dettmer** - Institute of Functional Genomics, University of Regensburg, Regensburg, Germany. shared first-authors (Antonia Hehemann, Jan Schueler). shared last-authors: Silke Haerteis, Katja Dettmer.

## Abstract

The chorioallantoic membrane (CAM) model represents a promising three-dimensional in vivo platform for preclinical drug testing in human tissues. In this study, we investigated whether the tissue penetration and distribution of benzbromarone, a known inhibitor of the Ca^2+^ activated chloride channel TMEM16A and potential therapeutic agent for autosomal dominant polycystic kidney disease (ADPKD), can be successfully visualized in human renal cyst tissue cultured on the CAM. To this end, desorption electrospray ionization mass spectrometry imaging (DESI-MSI) combined with an ultrahigh-resolution time-of-flight mass spectrometer was employed. We achieved spatially resolved molecular mapping of endogenous metabolites and lipids as well as the applied compound.

MSI enabled clear differentiation between CAM and cystic tissue based on their distinct lipid profiles. Benzbromarone was reproducibly detected in the cyst specimens and exhibited selective accumulation along the cyst epithelium, which is considered the principal site of action. These observations were complemented by multivariate analyses including Uniform Manifold Approximation and Projection (UMAP), and sparse multinomial logistic zero-sum classification. The data-driven approach confirmed molecular differences between tissue types and allowed accurate classification of drug-treated and untreated regions.

This study demonstrates that topically applied benzbromarone penetrates human renal cyst tissue in the CAM model and localizes to pharmacologically relevant tissue regions, notably the location of the Ca^2+^ activated chloride channel TMEM16A in the epithelial lining. The integration of high-resolution DESI-MSI with advanced statistical analysis provides a robust and label-free method to study drug distribution in human tissue grafts. Our findings contribute to the advancement of translational research in analytical chemistry and pharmacology.

## INTRODUCTION

Autosomal dominant polycystic kidney disease (ADPKD) is the most common inherited kidney disorder and is characterized by progressive cyst formation and enlargement, ultimately leading to loss of renal function.^1,5,2–4^ Despite advances in understanding disease pathogenesis, therapeutic options remain limited and approximately half of affected individuals progress to end-stage kidney disease by the age of 60 years.^6^ Renal cyst growth is promoted by Cl^-^ secretion and fluid transport into the cyst lumina which is significantly mediated by the Ca^2+^ activated Cl^-^ channel TMEM16A^7^. The benzofuran derivative benzbromarone belongs to the group of uricosurics and is an alternative drug for the treatment of hyperuricemia^8^. Benzbromarone inhibits not only the reabsorption of urate by URAT1 in the renal proximal tubule^9^ but also the Ca^2+^ activated Cl^-^transport by TMEM16A in a variety of tissues^10^. Remarkably, benzbromarone diminishes renal cyst growth *in vitro* and murine ADPKD models^7^. However, the efficacy in humans remains to be investigated.

The chorioallantoic membrane (CAM) model, a 3D in vivo model initially established for angiogenic and oncological research, enables the treatment and analysis of human renal cystic tissue obtained from nephrectomies^11^. The CAM is a well-vascularized extraembryonic membrane of fertilized chicken eggs localized next to the eggshell. Since complete immunocompetence is absent until day 18 of embryonic development, the CAM model allows the engraftment and investigation of various cells and tissues from different species for approximately one week^12^. Recently, we could demonstrate that blood vessels of the CAM and blood vessels from engrafted human renal ADPKD formed functional anastomoses within 12 hours after engraftment^13^. As the CAM model allows drug application, this model represents a potential platform for the investigation of novel therapies in ADPKD^14^. However, the spatial distribution of applied substances has not been analyzed previously. Mass spectrometry imaging (MSI) allows for the simultaneous determination of the spatial distribution of pharmaceuticals, their metabolites as well as endogenous chemical species in tissue sections making it a suitable approach for mechanistic pharmacological and toxicological studies and the direct correlation of drug and metabolite distribution with histological and immunohistochemical findings^19,15,18,17,16^. The two main mass spectrometry imaging techniques applied for the study of drug distributions in tissues are either atmospheric pressure matrix-assisted laser desorption/ionization (AP-MALDI), without and with laser-induced post-ionization to overcome ion suppression effects^20^, or (nanospray) desorption electrospray ionization (DESI) for desorption and ionization of compounds from the surface of tissue sections. They are typically hyphenated to triple quadrupole or high-resolution time-of-flight mass spectrometers for the targeted and nontargeted analysis, respectively, of drugs and their metabolites. DESI is based on a pneumatically-assisted spray of electrically charged solvent droplets that is directed onto the sample surface, inducing desorption and ionization of analyte molecules^14^. The resulting ions are transferred to the mass analyzer via a heated transfer capillary^21^. In this work, a DESI source was coupled to an ultrahigh-resolution time of flight mass spectrometer (HR-TOFMS) with mass spectral resolution of up to 200,000 full width at half maximum (FWHM) to analyze the spatial distribution of topically applied benzbromarone in human renal cysts engrafted onto the CAM in order to study its tissue penetration and distribution. Furthermore, machine learning yielded a sparse zero-sum LASSO mass spectral signature that clearly distinguished CAM and renal cyst tissue with and without benzbromarone.

## METHODS AND MATERIALS

The workflow used in this project is depicted in Figure 1 and details are described here:

**Figure 1.**
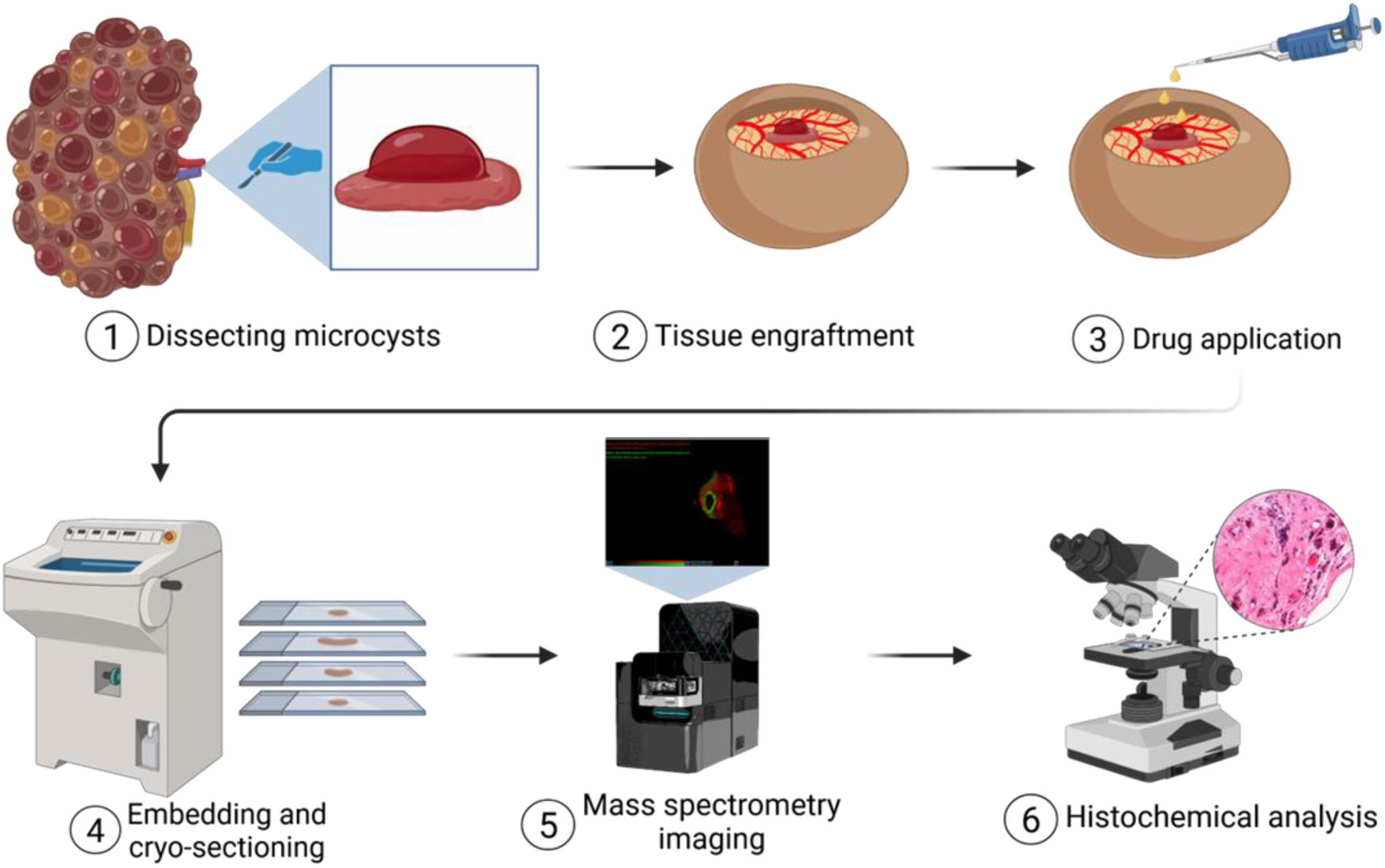
Workflow scheme.

### Chemicals

LC-MS grade methanol was purchased from VWR (Ismaning, Germany), polyvinylpyrrolidone (PVP) and benzbromarone from Sigma Aldrich (Steinheim, Germany), hydroxypropylmethylcellulose (HPMC) from Fisher Scientific (Schwerte, Germany), and leucine enkephalin from Waters (Milford, MA, USA). Water purified by the PURELAB Plus system (ELGA, LabWater, Celle, Germany) was used for all aqueous solutions. A stock solution of benzbromarone (50 mM) was prepared in DMSO and further dissolved in sterile isotonic 0.9% (w/v) sodium chloride solution to a final concentration of 50 µM.

### Tissue samples

ADPKD tissue was obtained from patients undergoing nephrectomy at the University Hospital of Regensburg, the St. Elisabeth Hospital Straubing, and the University Hospital Erlangen. The ethics committee of the University of Regensburg approved the experiment (no. 20-1886-101). All patients gave their written consent. After nephrectomy, the tissue samples were kept in MACS Tissue Storage Solution (Milteny, Germany). Microcysts with a diameter of approximately 1-2 mm and surrounding tissue were prepared from renal cyst tissue and engrafted onto the CAM as recently described^11^. In total 10 cysts from 4 patients were investigated (supplementary Table S1). The underlying genetic variants of the ADPKD patients were not determined. Cysts were engrafted on the CAM of individual eggs. Pork liver for quality control measurements was obtained from a local supplier.

### The CAM model

The CAM model was performed as described previously^11,13^. Briefly, fertilized eggs were incubated for 4 days in the ProCon incubator (Grumbach, Asslar, Germany). Then, a window was cut into the eggshell and sealed with Leukosilk^®^ (BSN medical GmbH, Hamburg, Germany). Microcyst tissue was engrafted onto the CAM on day seven of embryonal development. The tissue was positioned in the center of an agarose ring on the CAM to stabilize its position in the center of the eggshell window. Vitality of the eggs was checked daily, and macroscopic images were taken with a microscope (Leica^®^ M205A microscope). Fifty µL of benzbromarone (50 µM in sterile, isotonic 0.9% sodium chloride solution) were applied topically onto the cysts once a day for seven days. One hour after the last topical application, the tissue was removed and snap frozen in liquid nitrogen. Cysts 9 and 10 from patient 4 were additionally washed with PBS to investigate the effects of extra washing on subsequent drug measurements.

### Cryosections

Cryosections were prepared by means of a Leica CM3050 S cryostat (Leica Biosystems, Nussloch, Germany). A hydrogel containing 7.5% HPMC and 2.5% PVP was used as an embedding medium^22^. Medium was placed on the sample holder and frozen. Then, the frozen tissue was placed on top of it and fully covered with additional hydrogel. The tissue was sectioned at a thickness of 10 µm, before two consecutive tissue sections were thaw mounted onto a microscope slide. Between slides, 200 µm of tissue were trimmed. Sectioning for cysts used for quantification deviated from this routine as described below.

### DESI-HR-TOFMS analysis

All DESI-HR-TOFMS measurements were performed with a SELECT SERIES Multi Reflecting Time-of-Flight (MRT) mass spectrometer (Waters, Milford, MA, USA) equipped with a DESI XS source (Waters). An 11 elite syringe pump (Harvard Apparatus, Holliston, MA, USA) was used to deliver the spray solution. The instrument was set to operate mode for at least an hour prior to mass calibration using polyalanine (5 mg/mL polyalanine in 95 % methanol, 5 % water pipetted onto microscope slides). Mass spectral resolution of 180,000 was routinely achieved.

The desorption solvent was methanol/water (95/5 v/v) containing 100 pg/µL leucine enkephalin. The solvent flow rate was 2 µL/min. Pork liver measurements were executed daily as an instrument quality control. All measurements were performed in negative ion mode. The step size was 50 µm for ADPKD tissue and the scan time was 0.5 s. The capillary voltage was 0.7 kV and the cone voltage 20 V. The inlet tube was heated to 150 °C and the source temperature was 100 °C. The sprayer contact angle was 75 °. The nebulizing nitrogen gas pressure was 1 bar. Before measurements, sections were dried in a vacuum desiccator for 5 min.

### Benzbromarone quantification

For quantification of benzbromarone the kidney cysts were cryosectioned as follows. A 10-µm trim section was placed in 100 µL 80% methanol (80% MeOH; methanol/water, 80/20(v/v)). The next two consecutive 10-µm sections were thaw mounted onto a microscope slide. The subsequent 10-µm trim section was again transferred into 100 µL 80% MeOH. Then, 180-µm of the tissue was trimmed before the next 10 µm section was used for extraction. The extraction and measurement of the kidney cyst sections is described in the supplement.

The DESI measurement of the kidney cyst sections was performed as described above. Afterwards the data was lock-mass calibrated using leucine enkephalin (*m/z* 554.2620 in negative ion mode) and then processed in HDI (Version 1.6, Waters). A mass range of 50-2000 *m/z* and a *m/z* window of 0.005 Da were used and the 3,000 most intense peaks over the whole measurement were selected. The location of the benzbromarone signal on the cystic tissue was evaluated. Using a 1-µL syringe, 0.1 µL of a mixture containing 1 µmol/L benzbromarone and fluoresceine 25 µmol/L in 50% methanol was spiked on the consecutive tissue section. The spiked section was then measured and semi-quantitative benzbromarone quantification was performed following a standard addition approach as described in the results sections.

### Hematoxylin/Eosin staining

Slides were subjected to Hematoxylin/Eosin (H/E) staining after DESI-HRTOFMS measurements. The cryo-sections were first fixed with 4 % paraformaldehyde (PFA) in PBS (pH 7.4) for 10 min, followed by three washing steps with PBS (pH 7.4) for 5 min each, stained with H&E according to the standard protocol^11^, and dehydrated using an ascending alcohol series. Finally, cover slips were placed and mounted with DPEX medium.

### Data analysis

The imaging data was lock-mass corrected using leucine enkephalin (*m/z* 554.2620, negative ion mode) with the threshold set to 100 counts and a mass tolerance of 500 ppm. Initial data processing was performed using HDI (Version 1.6, Waters) with a mass range of 50 to 2000 Da, a *m/z* window of 0.005 Da, and selection of the 3000 most intense peaks. The resolution was determined by extracting a spectrum from the chromatogram and using the rescal tool (Version 1.0.1.0, Waters). The data was TIC normalized for visualization. HDI was also used to define regions of interest (ROIs) for CAM, cyst tissue (CT) and cyst tissue with benzbromarone (CTB).The annotated HE stain was used as a guide to correctly differentiate between CAM and CT as well as the benzbromarone signal to differentiate between CT and CTB when defining the ROIs. For cyst 1A, three different ROIs each were selected for the three regions resulting in 9 ROIs for this tissue section. The scan numbers included in the respective ROI were exported as vector and used to extract the respective data in R.

Further data analysis was performed in R (version 4.3.3)^23^ with RStudio (version 2023.12.2 Build 402) using the packages ggrepel (version 0.9.6)^24^, openxlsx (version 4.2.8)^25^, readxl (version 1.4.5) ^26^, tidyverse (version 2.0.0)^27^, data.table^28^ (version 1.17.2), fuzzyjoin^29^ (version 0.1.6) umap (0.2.10.0)^30^, and zeroSum (version 2.0.7)^31^. The data analysis strategy was developed entirely by the authors. AI (ChatGPT, OpenAI) was used solely as a supportive tool for generating and debugging R code. The generated code was critically reviewed, tested and validated.

For data preprocessing an inclusion list containing masses to be kept regardless of the cleanup steps (m/z = 554.262 (leucine enkephalin), m/z = 255.233 (palmitic acid, C16:0), m/z = 283.2643 (stearic acid, C18:0), m/z = 279.233 (linoleic acid, C18:2), m/z = 277.2173 (α-linoleic acid, C18:3), m/z = 303.233 (arachidonic acid, FA 20:4) as well as a list containing masses to be used to differentiate between tissue and background pixel (m/z = 885.5497 (PI 38:4), m/z = 766.5391 (PE 38:4), m/z = 572.4814, (Cer d34:1)) were defined. Based on the three masses in the latter list, pixels were divided into background and tissue pixels. The data was filtered to only keep masses with intensities that were nonzero in at least 20 tissue pixels. To remove noise and background, the ratio of the mean intensity in the tissue to the mean intensity in the background was calculated for each m/z value. Only masses with a ratio bigger than 1.5 were kept. ROIs determined in HDI are used to assign pixels to the CAM region, cyst region (CT) or cyst region with benzbromarone (CTB). Only pixels containing benzbromarone (m/z = 422.906) are kept in CTB and only pixels without benzbromarone are kept in CT. Missing m/z values in each m/z column were imputed using the smallest value in the respective column divided by a randomly selected number between 5 and 5.5. All intensities were log2-transformed.

For further analysis, 17 cyst data sets were analyzed together. A data set includes three ROIs defined for the different tissue regions. For cyst 1A three different ROIs were set for each region resulting in 3 data sets for this section. Data from cysts 4 and 5 were not included in the analysis as no CAM region was visible. To align m/z values across datasets, all m/z values were compiled and the difference between neighboring m/z values calculated. The m/z values were then sorted into bins with a tolerance of 3 mDa. The average m/z of each bin was determined. Each cyst data set was matched to the corresponding bin IDs. For each pixel in a file that belonged to the same ROI and bin ID, the mean intensity was calculated. All bins were combined with the average bin mass added. For further analysis only columns with a maximum of 25% NA values were used. Missing values were then imputed using the smallest value in the column divided by a random number between 5 and 5.5. The data was log2 transformed and standardized using z-score transformation. A UMAP projection was generated using n-neighbors = 40. No further normalization was performed.

For classification of tissue regions, a multinomial logistic regression model with the Least Absolute Shrinkage and Selection operator (LASSO) to enforce sparsity and a zero-sum constraint to obtain normalization invariant signatures was fitted to the log2 transformed data (R package zeroSum, version 2.0.7^31^). to enforce LASSO regression the elastic net parameter α was set to 1. For model training, highly correlated features (r >0.9) were removed and only the feature with the higher intensity was kept. Two data sets stemming from cyst 1A were combined and used for model training, the third data set and the data sets from the other sections were used for independent testing. To generate a robust and stable signature with reliable features a two-step procedure was conducted. In the first step, 100 repeated stratified resampling runs (70% training, 30% validation) were performed. In each run, a multinomial logistic zero-sum LASSO model with 10-fold cross-validation was trained. The selection frequency of all features across all runs was calculated to determine feature stability. Features with a selection frequency > 50% in at least one class were used to train in the second step the final model on the full training dataset, employing again 10-fold cross-validation to determine the optimal regularization parameter (λmin) that controls sparsity of the model. For the multinomial logistic LASSO regression model the class vector was set to 1 = CAM, 2 = CT, and 3 = CTB, respectively.

## RESULTS AND DISCUSSION

### Detection of benzbromarone

Benzbromarone was spotted on pork liver sections to establish analysis by DESI-HR-TOFMS. Pork liver was used as model tissue due to its homogeneity. Benzbromarone was detected in negative ion mode and the tune page parameters were optimized manually using the signal intensity of the [M-H]^-^ ion at *m/z* 422.906. A mass spectrum of benzbromarone spotted on a pork liver section is shown in supplementary Figure 1A. Decreasing concentrations of benzbromarone (50 µM – 20 nM, 0.6 µL) were spotted onto pork liver sections to establish the detection limit. Supplementary Figure S1 shows the spotted data using square root scaling (1B) and log scaling (1C) for the intensity scale. Looking at the log scaled data reveals smearing at higher concentrations, as the spotted benzbromarone is smeared toward the right side of the spot. Regions of interest (ROIs) covering 1405 scans (pixel size 50 x 50 µm) were set for each spotted level to extract the signal intensity of benzbromarone. The average intensity was plotted over the spiked concentration (supplementary Figure S1D). We were able to detect the spotted benzbromarone at a concentration as low as 0.08 µM. With a droplet of 0.6 µL covering an area of about 1,194 scans, this in 0.04 fmol/scan. We calculated a median spectrum using the ROI of this spotted level to determine the signal-to-noise ratio (SNR). Root Mean Square (RMS) noise was determined using all signals not exceeding 5 % of the maximum signal as noise. For benzbromarone this results in a SNR of 7 (*m/z* 422.906). However, the determined LOQ has to be considered with caution as the spot experiment does not account for differences in desorption and matrix effects experienced by benzbromarone spotted onto the tissue section versus benzbromarone located in the tissue.

Next, we analyzed tissue sections of microcysts cultivated on the CAM for seven days (see workflow, Figure 1). During cultivation on the CAM, cysts were topically treated with 50 µL of benzbromarone (50 µM) once per day. Several sections were prepared for each cyst-CAM sample. Sections of three different cysts from patient 1 and 4 are shown in Figure 2

**Figure 2.**
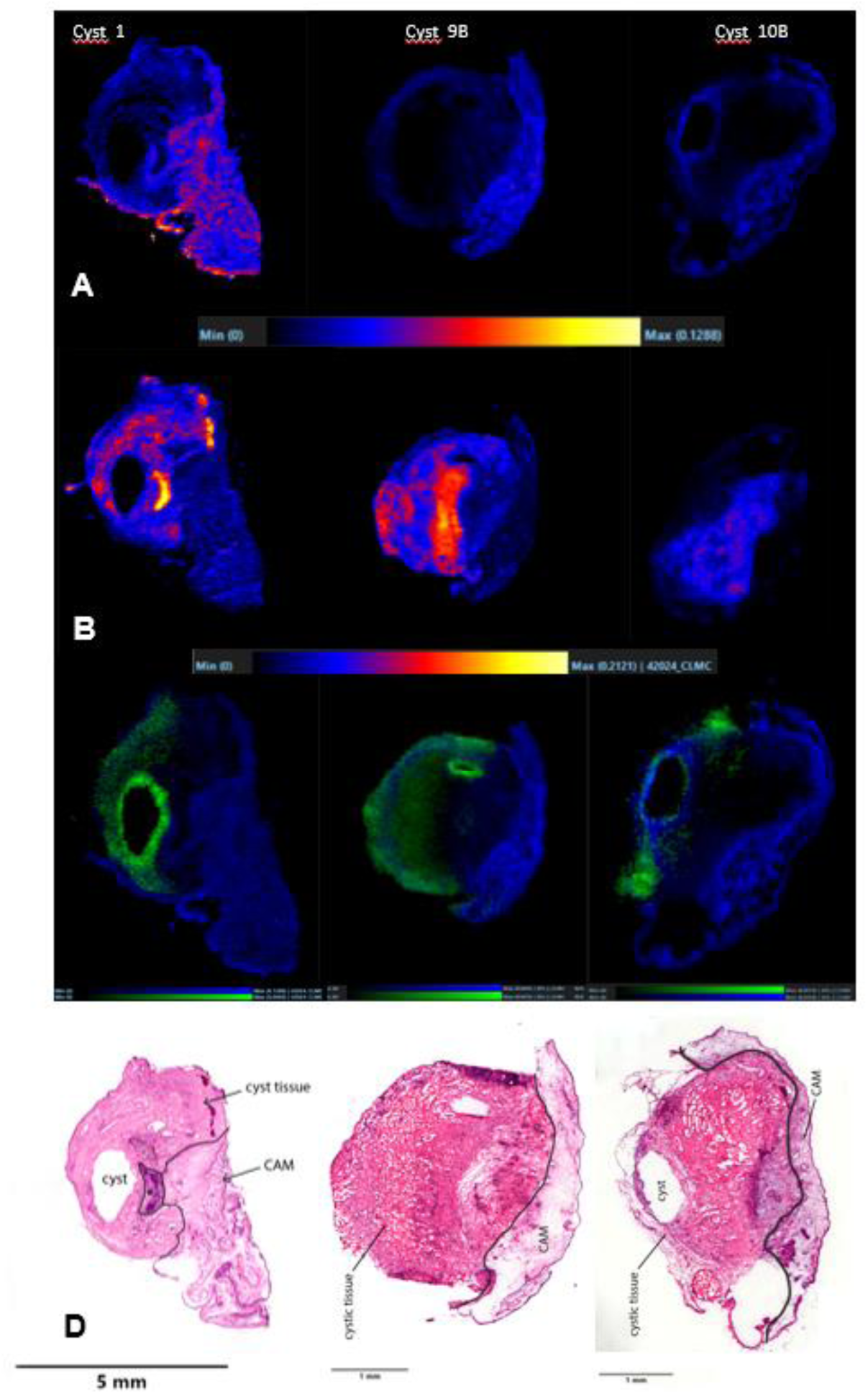
Ion and H/E images for three cyst-CAM specimens. A) TIC normalized signal of phosphatidylethanolamine PE (38:4) at *m/z* 766.539, [M-H]^-^. B) TIC normalized signal of ceramide (Cer d34:1) at *m/z* 572.481, [M+Cl]^-^. C) Overlay of the benzbromarone signal at *m/z* 422.906 ([M-H]^-^) and PE (38:4) to visualize the location of benzbromarone on the cyst-CAM tissue. Mass spectral images were acquired with a step size of 50 µm, a scan time of 0.5 s and a flow rate of 2 µL/min. D) Corresponding H/E staining. Slides were subjected to H/E staining after DESI-HR-MS analysis. * Obliterated vessel.

Figure 2A shows the ion image of a structural lipid PE 38:4 (m/z = 766.5391, C_43_H_78_NO_8_P, [M-H]^-^, −0.13 ppm) used to visualize CAM tissue, while Figure 2B shows the ion image of Cer d34:1 (m/z = 572.4814, C_34_H_67_NO_3_, [M+Cl]^-^, 0 ppm) used to visualize the cystic tissue. Figure 2C shows the ion images of benzbromarone (*m/z* 422.906) for the three cyst-CAM samples overlayed with PE38:4 to reveal the tissue distribution of benzbromarone. During embedding and sectioning the orientation of the tissue was precisely maintained and only lateral sections were produced. Therefore, the CAM was located on the right side of the section. The benzbromarone signals are detected more at the top of the cyst tissue (left side of the section). This is to be expected as the drug is dripped onto the cyst tissue during topical treatment and most of it accumulates on top and on the sides of the cyst tissue. This also explains the gradual loss of benzbromarone signal intensity towards the inside of the cyst tissue. Notably, for all three cysts an accumulation of benzbromarone around the cyst cavity is observed with higher signal intensities for the epithelial lining of the cyst, where it is expected to act as TMEM16A inhibitor, thereby reducing Cl^-^ secretion into the cyst^32,7^ Supplementary Figure S2 shows two cysts (cyst 4 and 5) from patient 2 treated topically with benzbromarone. Benzbromarone was clearly detected in both tissue specimens. Unfortunately, the CAM was removed during excision of the cyst tissue, therefore, information regarding the location of the tissues on the CAM had been lost. Notably, in cyst 4, a metabolite of benzbromarone, namely hydroxybenzbromarone (*m/z* 438.9008, C_17_H_12_Br_2_O_4_, [M-H]^-^, −0.23 ppm), was detected (Figure S2C). This metabolite was not observed in the other specimens analyzed. Supplementary Figure S3 depicts two cysts (cyst 6 and 7) from patient 3 including two sections from cyst 7. A clear distinction between CAM and CT can be observed based on the structural lipids PE 38:4 and Cer d34:1. As expected, the benzbromarone signal for cysts 6 and 7 was found at the top of the cystic tissue with a decreasing signal intensity towards the CAM. Similar benzbromarone distributions could also be seen in further cysts samples from patient 1 shown in Supplementary Figure S4. No cyst cavities were observed in either sample set. Cysts 9B and 10B from patient 4 are shown in Figure 2. These cysts were washed with PBS after explantation to remove any potential accumulation of benzbromarone on the outside of the tissue that could be spread over the tissue during cryosectioning. The washed cysts should only show a signal for benzbromarone that has actually penetrated the cystic tissue.

## Quantification of benzbromarone Spatially resolved quantification

Quantification by DESI-MS is still difficult to achieve. Spike-on experiments ignore both differences in desorption of endogenous and spiked compound as well as matrix effects. Adding internal standard to the desorption solvent disregards differences in desorption and matrix effects as well.^33^ Homogenizing the tissue with internal standard and measuring homogenate extracts leads to good quantification results, albeit at the loss of spatial information.^34^

Here, we pursued two strategies for spatially resolved benzbromarone quantification (see supplementary Figure S5) employing either LC-MS/MS analysis of tissue extracts (extraction method) or on-tissue standard addition (standard addition method). For testing and comparison of both quantification approaches a total of four consecutive 10-µm sections were prepared for each benzbromarone treated cyst analyzed. For the extraction method tissue sections 1 and 4 were extracted with 80% methanol/ *tert*-butyl-methylether (see supplementary methods). The second and third consecutive 10-µm sections were thaw mounted onto the same microscope slide. For the standard addition method, tissue section 2 was measured to elucidate the location of benzbromarone signals. Then, 0.1 µL of a mixture containing 1 µmol/L benzbromarone and fluoresceine 25 µmol/L in 50%methanol was spiked on the consecutive tissue section 3 in areas that contained benzbromarone. Three spike-on’s were executed per section. After MSI of the spiked section, 3 ROIs were selected in the locations of the spikes that included areas with added and without added benzbromarone. The data was filtered in R and separated into two data frames with one containing pixels with both a benzbromarone and a fluoresceine signal (spike signal) and one with pixels containing only benzbromarone (signal of the “endogenous” benzbromarone). The benzbromarone concentration was then calculated via standard addition. Analyzing the three ROIs with spikes leads to an average amount of 0.14 fmol benzbromarone per pixel for cyst 8 (patient 3). With a pixel size of 50 x 50 µm and a thickness of 10 µm this leads to a total volume of 25,000 µm^3^ or 25 pL. This results in an average estimated benzbromarone concentration of 5.5 µmol/L per pixel.

For the extraction method, the first and last tissue section were individually extracted and the total benzbromarone amount was determined by LC-MS/MS (supplementary Figure S5/supplementary methods). This total amount was used to estimate the amount of benzbromarone per pixel. The MSI data of the unspiked tissue section 2 was filtered in R for selection of pixels with a benzbromarone signal only. The total benzbromarone signal was calculated and for each pixel the percentage contribution to this total signal was determined. Using the data from the extracted sections, the average total amount of benzbromarone was calculated and equated to the total benzbromarone signal. From that the amount of benzbromarone in each pixel was calculated. This resulted in an average amount of 0.32 fmol benzbromarone per pixel for cyst 8A (patient 3). This leads to an average concentration of 12.8 µmol/L using the above-mentioned volume for a pixel. This two-fold quantification approach was performed for two cyst samples as summarized in Table 1. Furthermore, two additional sections per cyst were subjected to extraction and the subsequent section was imaged (see B samples, Table 1). Interestingly, absolute benzbromarone amounts across all extracted sections did not vary greatly (average amount 46.3 fmol, RSD 11.7%, n=4), but the number of pixels did (120 – 751), resulting in varying pixel concentrations.

**Table 1.**
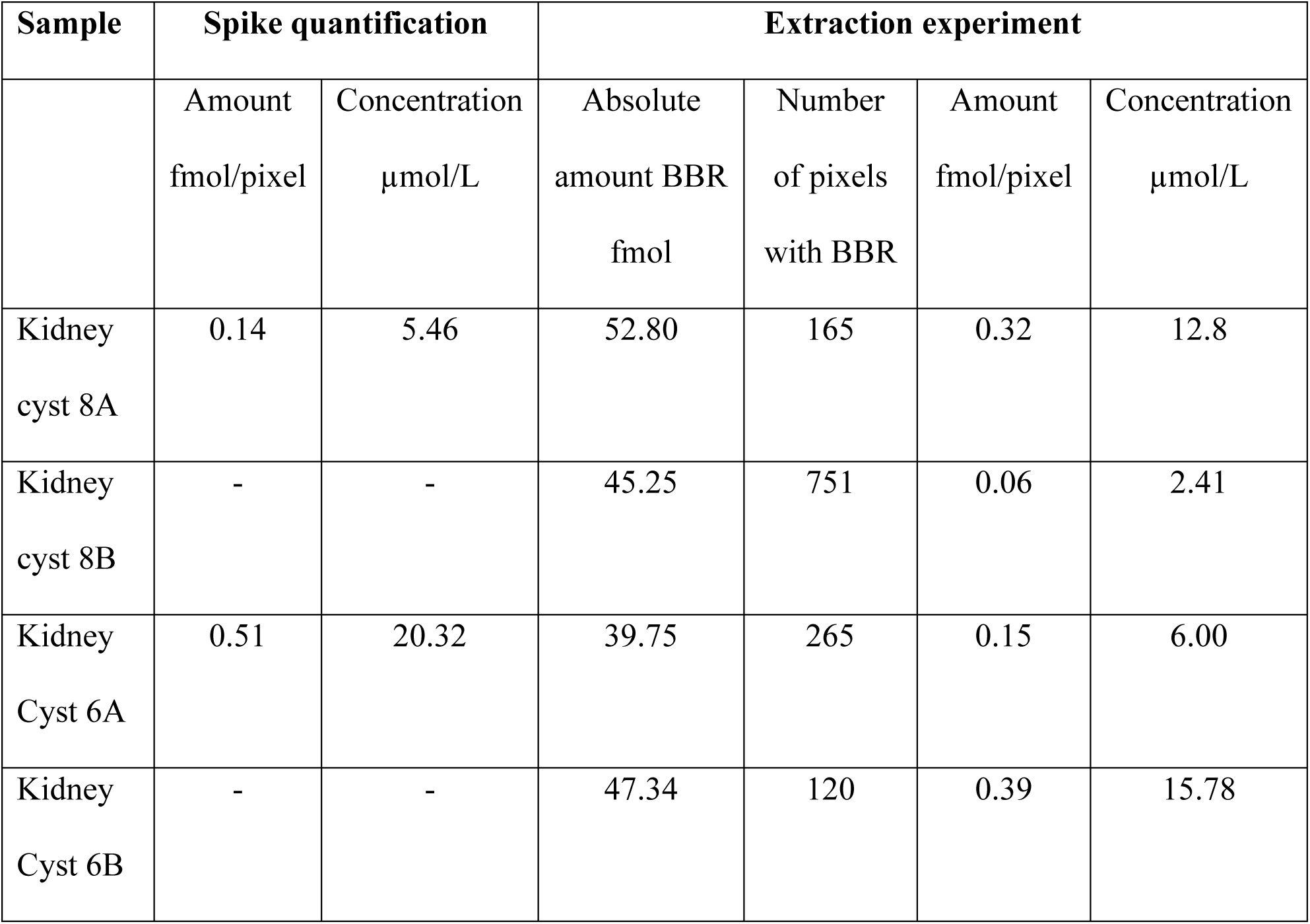
Results of the benzbromarone quantification for cysts 6 and 8 of patient 3). The volume of a pixel was calculated based on a the step size of 50 µm x 50 µm and a tissue cutting thickness of 10 µm corresponding to a volume of 25000 µm^3^ or 25 pL.

Benzbromarone has been shown to effectively block the flux of anions through the TMEM16A channel with a half maximum inhibitory concentration (IC_50_) of 9.97 µmol/L.^35^. A QPatch whole-cell electrophysiology screen yielded even lower IC_50_ values of 2.35±0.96 µmol/L^36^. More recently, using the human cystic fibrosis bronchial cell line CFBE41o-as a model, 1 µmol/L of benzbromarone was shown to inhibit TMEM16A effectively.^37^ Thus our estimated concentrations fall within the IC_50_ concentrations reported for benzbromarone against TMEM16A.

While for the spike, a section thickness of 10 µm was assumed, keep in mind that the desorption solvent does not fully penetrate the tissue resulting in incomplete analyte desorption. The spike experiment is also affected by the previously mentioned problems of ion suppression and altered desorption kinetics. As such, this method gives only an estimate of the benzbromarone concentration. Compared to the spike experiments, a higher benzbromarone concentration was found using the extraction method in cyst 8A, as complete extraction of the 10-µm section can be assumed. However, cyst 6A shows the reverse, with the concentration determined from the spike experiment being 3.5 times higher than the concentration from the extraction experiment. Further extraction experiments for cyst 6B and cyst 8B resulted in benzbromarone concentrations of 0.39 fmol/pixel (15.78 µmol/L) and 0.06 fmol/pixel (2.41 µmol/L). The concentrations determined via the spike experiment vary significantly. Between each measured section 220 µm of tissue were trimmed, which could cause the observed differences in concentration due to differences in tissue penetration of benzbromarone. The aforementioned drawbacks of the spike experiment could explain why we determined a higher concentration of benzbromarone for cyst 6A in the spike experiment. A further problem could be the correlation between fluorescein and benzbromarone. If fluorescein is not detected at each pixel where benzbromarone/fluorescein was spiked, the spike benzbromarone signal would be counted as an endogenous benzbromarone signal leading to over-quantification. Overall, the quantification approaches yield only an approximate estimate of benzbromarone tissue concentrations suggesting that these are in the low micromolar range.

### Visualization of feature space

As detailed in the data analysis section, 17 cyst data sets from three patients were combined. After alignment, the data were z-transformed on a per-dataset basis and subjected to Uniform Manifold Approximation and Projection (UMAP) analysis. UMAP showed a clear distinction between CAM and the cyst tissue in general but also between CT and CTB (Figure 3).

**Figure 3:**
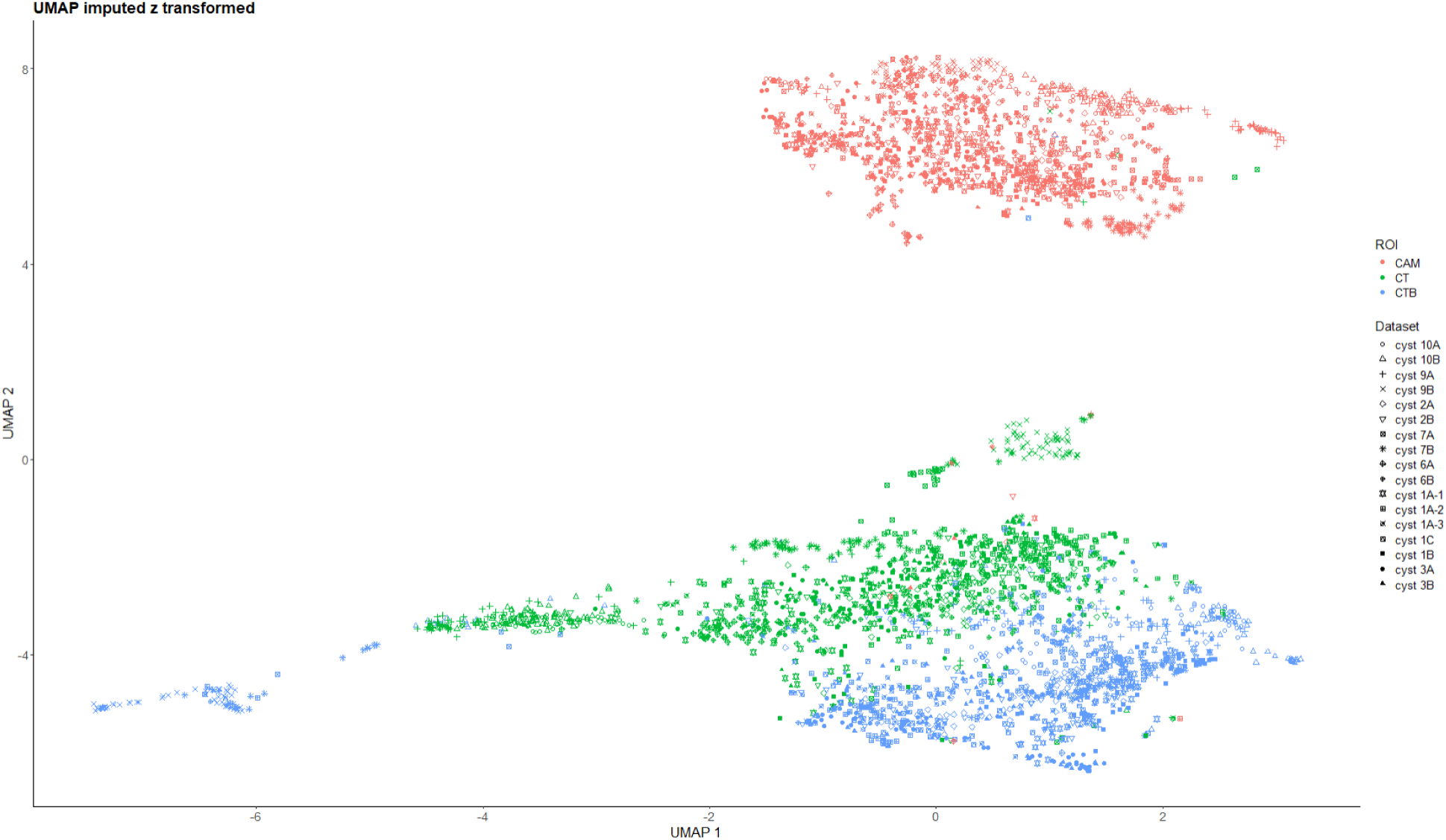
UMAP analysis of 17 cyst data sets from 3 patients. Datasets are visualized by different symbols and tissue regions by color. A clear distinction is seen between CAM and CT/CTB as well as CT and CTB.

### Classification of tissue regions

Next, we investigated whether it is possible to determine defined metabolic signatures that allow prediction of tissue types potentially reducing the need for additional histological and immunohistochemical stains for tissue classification. To this end, a multinominal logistic regression model with zero-sum constraint was fitted to the data. Least absolute shrinkage and selection operator (LASSO) regularization was used as it enables feature (masses) selection and coefficient shrinkage.^40,39,38^ It results in a sparse model with a low number of non-zero coefficients. The zero-sum constraint enables reference point insensitive data analysis obviating the need for prior normalization of the data^31^. For each data set, ROIs were set on CAM tissue, CT tissue and CTB tissue. To train a model, two ROI data sets of cyst 1A were used (Supplementary Table S3). The classes were set at 1 = CAM, 2 = CT and 3 = CTB, respectively. The non-zero coefficients of the models using λ min are given in Table S2. To predict CAM, 9 masses are needed, 20 masses for prediction of CT, and 8 masses for prediction of CTB. The signatures were then used to classify CAM, CT, and CTB tissue areas in sections of 15 cyst data sets (Supplementary Table S3). Figure 4 shows the classification for cyst 1B, 2A and 3A. It should be noted that training was done on data from cyst 1A while classification was performed on a consecutive section, 1B. Clear classification of CAM, CT and CTB is regularly achieved. The confusion matrices for cysts 1B, 2A and 3A are shown in Figure 4 as well. 100 % of the CAM pixels, 74.29 % of CT pixels and 100 % of CTB pixels were correctly classified for cyst 1B with only CT pixels being misidentified for CAM pixels with 10 % and CTB pixels with 15.71 %. It should be noted that the CTB ROIs are defined by the benzbromarone signal (see Methods and Materials, Data Analysis) and that benzbromarone (m/z = 422.906) is one of the features used to classifiy the CTB regions. As such the 100% correct classification of CTB is not surprising. Classification for cyst 1B achieved over all pixels an accuracy of 91.43 %, for cyst 2A an accuracy of 66.51 % and for cyst 3A an accuracy of 94.71 %. Classification was then applied to data from patients 1, 3 and 4. Supplementary Figures S6-S17 show the classification results for cyst 1A/C, 2B, 3 B, 6 A/B, 7A/B, 9 A/B and 10 A/B. Clear classification of CAM and CTB is achieved here as well. CTB regions were always classified correctly at 100%. This is expected, as CTB pixels were required to contain m/z 422.906, the major ion of benzbromarone, which also exhibited the absolute coefficient (−0,78). CAM was, in most cases, correctly classified at 87-100%, which is attributed to the distinct lipid composition of this tissue. The highest misclassification rate was observed for CT. This may be explained by patient-to-patient variability, as well as gradients in nutrient supply related to tissue vascularization and ROI selection with respect to these gradients.

**Figure 4.**
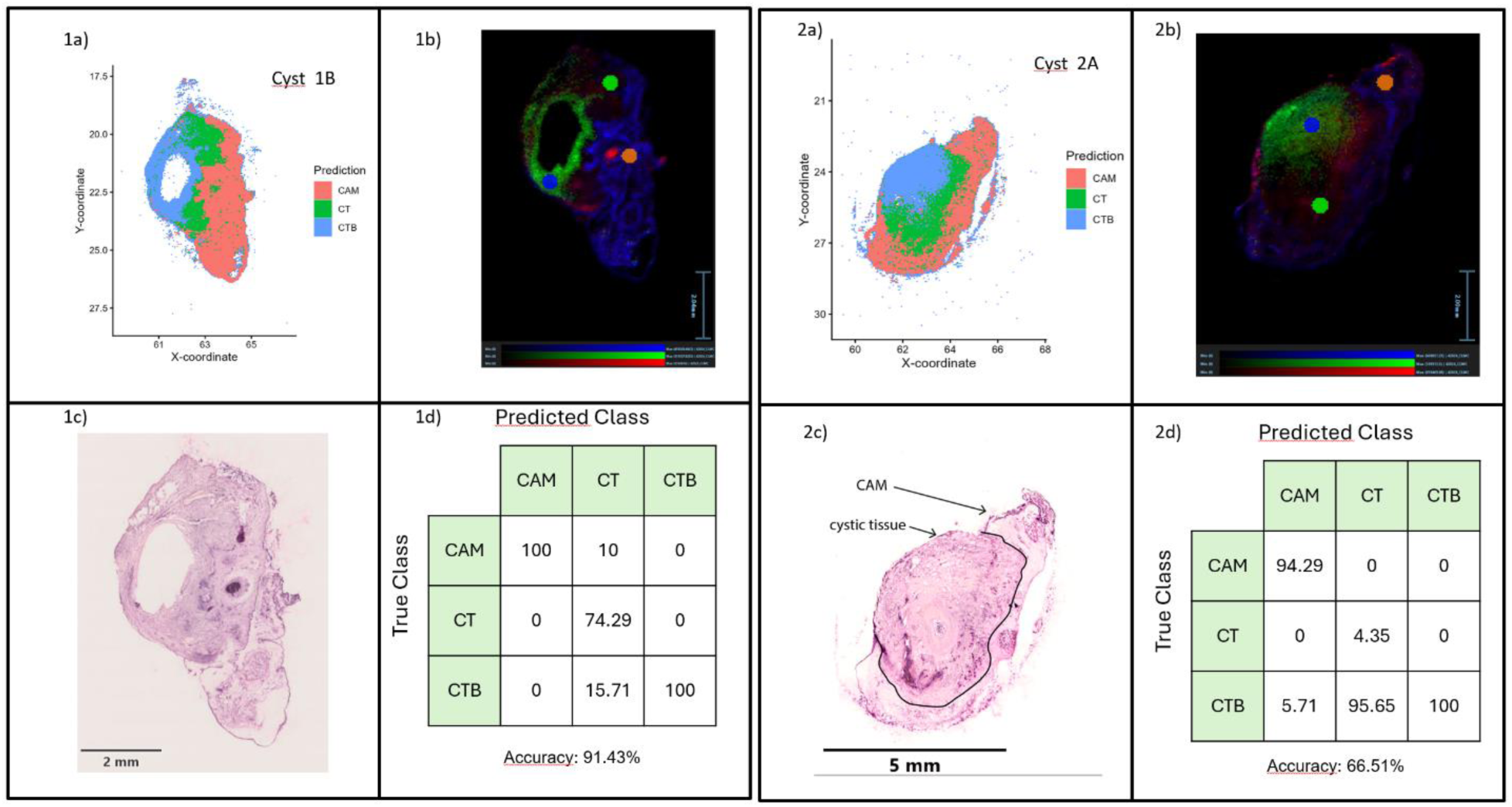
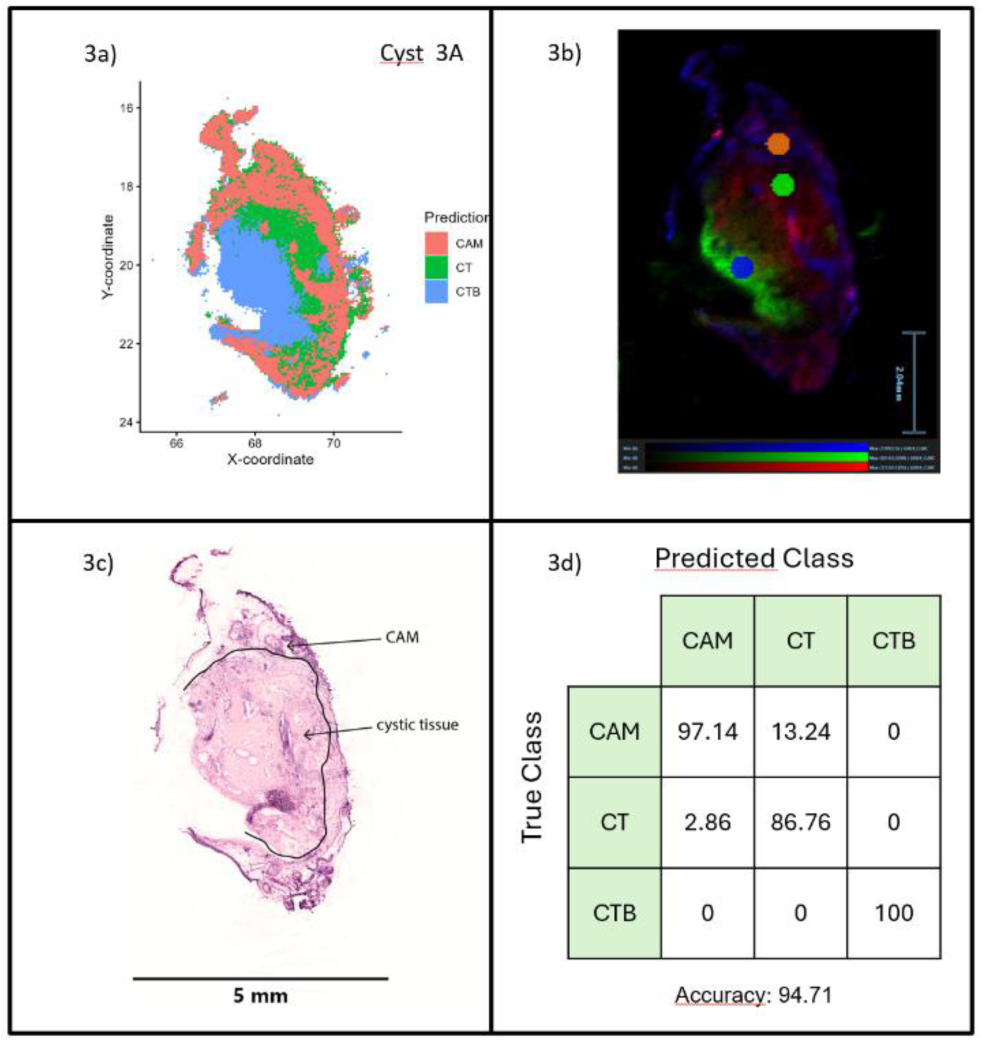
Results of the classification experiments for cysts 1B, 2A and 3A of patient 1. a) classification of CAM, CT, and CTB tissue areas for three cyst specimens using LASSO zero-sum signatures (1 = CAM, 2 = CT, 3 = CTB), b) an overlay of three signals (m/z = 422.906 (benzbromarone, green), m/z = 572.4824 (Cer d34:1, red), m/z = 766.5391 (PE38:4, blue) with the ROIs used for data analysis shown for each cyst, c) the HE stain of each cyst and d) the confusion matrix and the the classification accuracy for each cyst. It should be noted that prediction for cyst 1 was performed for a consecutive section (labelled 1B) that was approximately 860 µm away from the section used for training.

## CONCLUSIONS

DESI-HR-TOFMS imaging enabled direct visualization of benzbromarone distribution in human ADPKD cyst tissue maintained on the CAM. After topical application, benzbromarone was consistently detectable within cystic tissue and showed preferential localization at the cyst epithelium, where TMEM16A-mediated chloride secretion is considered to contribute to cyst expansion. Estimates obtained by on-tissue standard addition and extract-based quantification placed local benzbromarone levels in the low micromolar range, which is compatible with previously reported TMEM16A inhibitory concentrations.

The same imaging data also separated CAM, cyst tissue, and benzbromarone-containing cyst regions on the basis of their molecular profiles. UMAP and sparse zero-sum multinomial LASSO modelling confirmed this separation across additional cyst datasets. Because the benzbromarone ion itself contributed to the definition and classification of drug-containing regions, these classification results should be read as supportive of spatial drug mapping rather than as an independent tissue signature. This work therefore provides proof of principle that human renal cyst tissue on the CAM can be coupled with high-resolution mass spectrometry imaging to study local drug access in a preserved tissue context. The data show that a TMEM16A inhibitor can reach the epithelial compartment of human ADPKD cyst tissue, providing a basis for subsequent studies that combine spatial drug mapping with functional readouts

## Limitation

While our study provides compelling evidence for the applicability of DESI-MSI to visualize drug distribution in human cystic tissue on the CAM model, certain limitations should be acknowledged. First, the number of biological replicates was limited, with analyses based on ten cyst samples from four patients, the mutational status of which had not been available. Although the data are internally consistent, broader validation across a larger cohort would strengthen the conclusions. Second, only a single compound, benzbromarone, was investigated. The generalizability of the CAM-DESI-MSI platform to other drug classes remains to be explored. Third, while we focused on topical application, future studies should assess intravascular application for systemic distribution, which are feasible within the CAM model. These considerations will be addressed in subsequent investigations building upon our initial proof of concept.

## ASSOCIATED CONTENT

## Supporting Information

### Supplementary Tables

Supplementary Table S1: Samples analyzed. 17 sections stemming from 10 cysts and 4 patients were measured in total.

Supplementary Table S2: Non-zero coefficients of the multinomial zero-sum LASSO model.

### Supplementary Figures

Supplementary Figure S1: Optimization of benzbromarone detection.

Supplementary Figure S2: Ion images for two cyst/CAM specimens from patient 2.

Supplementary Figure S3: HE and Ion images from two additional cyst specimens obtained from patient 3.

Supplementary Figure S4: Ion images from two washed cysts obtained from patient 4.

Supplementary Figure S5: Workflow employed for benzbromarone quantification.

Supplementary Figure S6: Results of the classification experiments for cysts 1A for patient 1

Supplementary Figure S7: Results of the classification experiments for cysts 1C for patient 1

Supplementary Figure S8: Results of the classification experiments for cysts 2B for patient 1.

Supplementary Figure S9: Results of the classification experiments for cysts 3B for patient 1

Supplementary Figure S10: Results of the classification experiments for cysts 6A for patient 3

Supplementary Figure S11: Results of the classification experiments for cysts 6B for patient 3

Supplementary Figure S12: Results of the classification experiments for cysts 7A for patient 3

Supplementary Figure S13: Results of the classification experiments for cysts 7B for patient 3

Supplementary Figure S14: Results of the classification experiments for cysts 8A for patient 4

Supplementary Figure S15: Results of the classification experiments for cysts 8B for patient 4

Supplementary Figure S16: Results of the classification experiments for cysts 9A for patient 4

Supplementary Figure S17: Results of the classification experiments for cysts 9B for patient 4. Supplementary Methods File

## Author Contributions

All authors contributed to the drafting and revising of the manuscript, and approved unanimously to its final accepted version # Authors should be considered as shared first-authors. * Authors should be considered as shared last-authors.

## Declarations

### Ethics approval and consent to participate

Informed consent was obtained from all subjects and/or their legal guardians and experiments had been approved by the ethics committees of the University of Regensburg (no. 20-1886-101) and the Friedrich-Alexander-University Erlangen-Nuremberg (no. 23-404-Bn). All patient data was encrypted, and every patient was assigned an internal patient ID. All data protection guidelines from the University of Regensburg were met.

## NOTES

The authors have no conflict of interest to disclose.

