## Supplementary material for "DESI-MS-Based Analysis of Drug Distribution in Human Renal Cystic Tissue Using the Chorioallantoic Membrane (CAM) as a 3D In Vivo Model"

### SUPPLEMENTARY METHOD

#### Extraction and LC-MS/MS analysis of tissue sections

The 10-µm sections in 80% MeOH were vortexed and stored at -80°C until further analysis, but at least overnight. For further extraction, the samples were brought to room temperature, vortexed and centrifuged at 10,000×g at 4°C for 5 min. The supernatant was passed through 3kDa cut off filters (VWR, Darmstadt, Germany) to remove the embedding medium. The filters were washed twice with 80% MeOH before the extract was transferred onto the filter and centrifuged at 150 g, 4°C for 5 minutes. The tissue pellet was washed with 50 µL 100% MeOH. Upon centrifugation at 10,000×g at 4°C for 5 min the wash was also transferred onto the filter and centrifuged. The filtered extracts were dried down using an infrared vortex vacuum evaporator (CombiDancer, Hettich AG, Baech, Switzerland). To ensure complete extraction of benzbromarone the tissue pellet was washed twice with 50 µL *tert*-butyl-methylether (MTBE). The MTBE extracts were combined with the previous extracts and the solvent was evaporated under a gentle stream of nitrogen. The residue was resuspended in 100 µL acetonitrile. The extracts were measured on a TripleQuad 6500+ mass spectrometer (Sciex, Framingham, Massachusetts, USA) coupled to an ExionLC™ 30AD HPLC/UHPLC system (Shimadzu, Kyoto, Japan). An AQUITY Permier HSS T3 C18 column (1.8 µm, 2.1 µm x 100 µm, Waters) with a VanGuard pre column (1.7 µm, 2.1 µm x 5 mm) was used. The flow rate was set to 0.3 mL/min and the oven temperature was set to 35°C. Mobile phase A was water with 0.1 % formic acid and B was acetonitrile with 0.1 % formic acid. Gradient elution started at 60%B and went up to a 100 % B in 6 minutes. It was then kept at a 100 % B for 1 minute before the eluent composition went down to 60 % B again to equilibrate for 3 minutes. 2 µL of sample were injected. Electrospray ionization in negative ion mode and multiple reaction monitoring (MRM)-mode was used. The ion spray voltage was kept at 4.5 kV and the source temperature at 400 °C. Ion source gas 1 and 2 were set to 60 psi, the curtain gas to 45 psi and the collision gas to 9 psi. Four transitions were monitored for benzbromarone that are listed in table 1. The clustering potential for each transition was 100 V, the entrance potential 10 °V and the dwell time 10 ms.

*Table 1: Monitored transitions of benzbromarone in the MRM mode with their corresponding collision energy.*

| Analyte | Molecular Weight [Da] | Q1 mass [Da] | Q3 mass [Da] | CE [V] |
|---|---|---|---|---|
| Benzbromarone 1 | 424.1 | 422.9 | 78.8 | 100 |
| Benzbromarone 2 | 424.1 | 422.9 | 80.8 | 100 |
| Benzbromarone 3 | 424.1 | 422.9 | 250.8 | 41 |
| Benzbromarone 4 | 424.1 | 422.9 | 406.9 | 46 |

Benzbromarone quantification was performed by standard addition. After the initial measurement, four aliquots of 20 µL were taken from the extract and dried down. The residue was resuspended in 20 µL acetonitrile with different benzbromarone concentrations (no benzbromarone, 1 nmol/L benzbromarone, 3 nmol/L benzbromarone and 5 nmol/L benzbromarone). The extracts were then measured as described above.

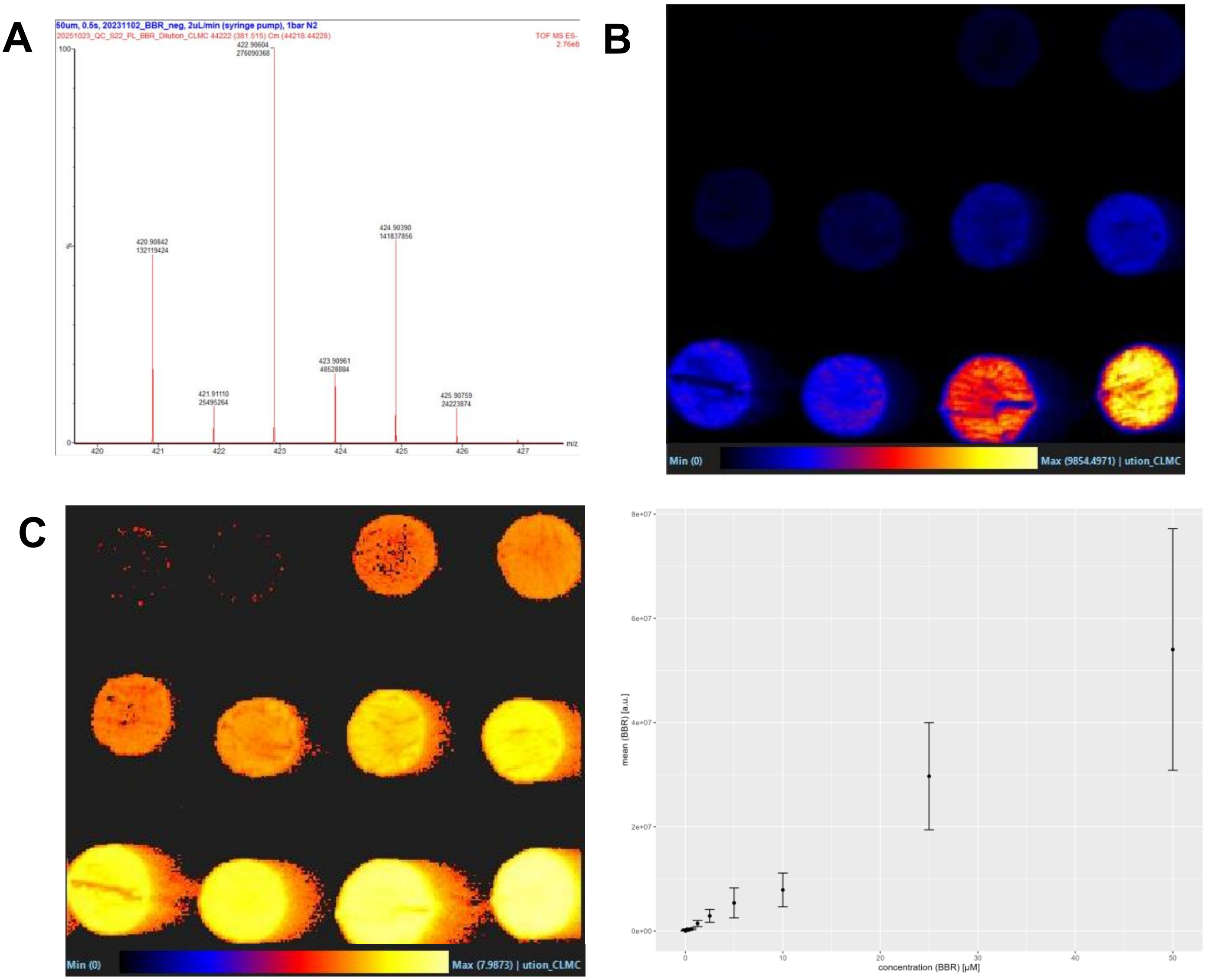


Supplementary Figure S1: Optimization of benzbromarone detection.
A) Benzbromarone was spotted on pork liver sections in concentrations of 50 µmol/L, 25 µmol/L, 10 µmol/L, 5 µmol/L, 2.5 µmol/L, 1.25 µmol/L, 0.63 µmol/L, 0.31 µmol/L, 0.16 µmol/L, 0.08 µmol/L, 0.04 µmol/L and 0.02 µmol/L (bottom right corner to top left corner). Analysis was performed in negative ion mode with a step size of 50 µm and a scan time of 0.5 s. The signal at m/z = 422.9059 ($[M-H]^-$) is shown. Square root scaling was applied.
B) Log scaling was applied to data shown in A. For high spike levels a wash-out is observed. This is attributed to the high amounts of benzbromarone applied on the tissue.
C) ROIs (average 1300 pixel) were set for each spike level and plotted against the concentration. Mean intensity ± standard deviation is shown.

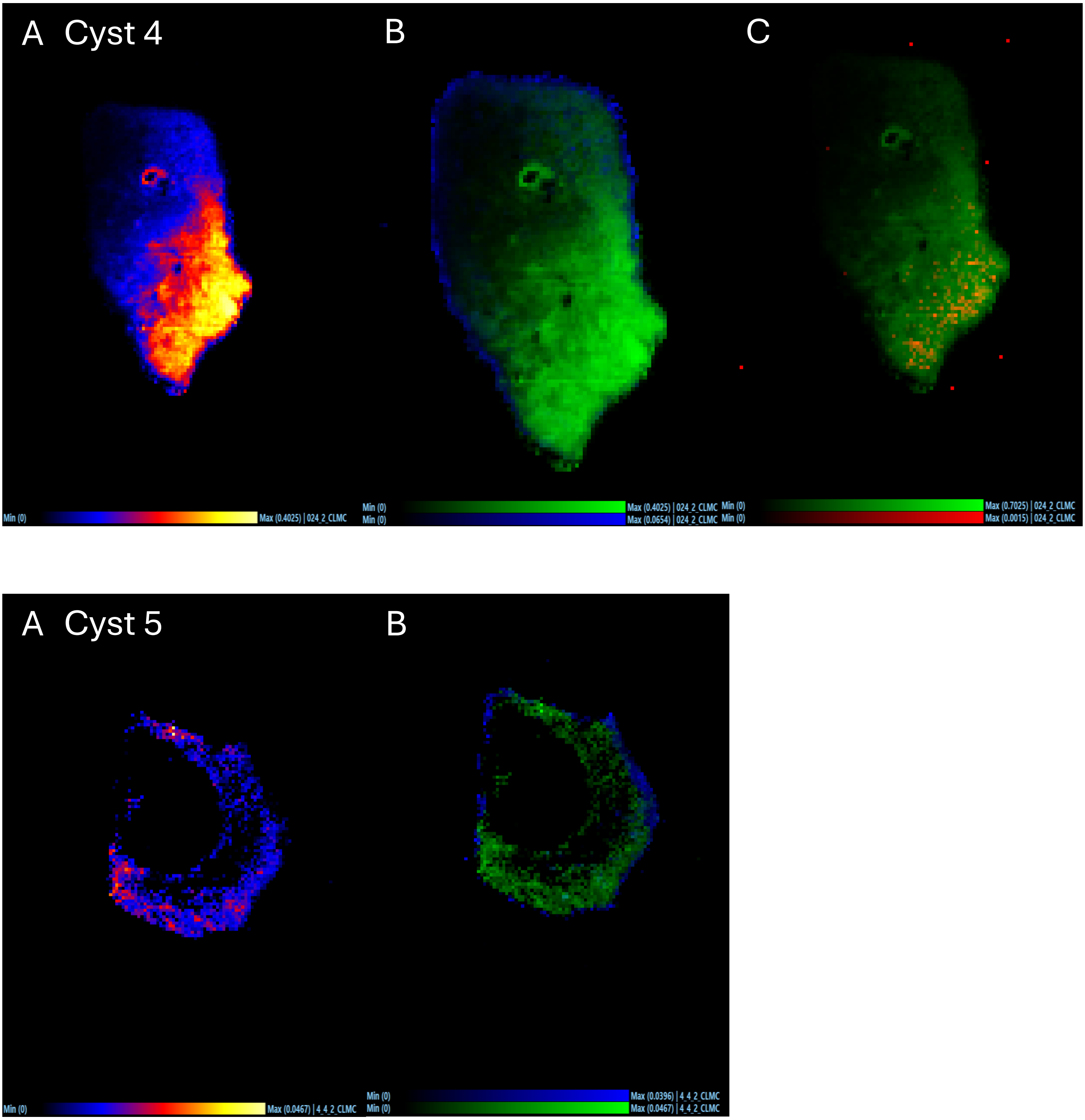


Supplementary Figure S2: Ion images for two cyst/CAM samples from patient 2. Upper row shows ion images of cyst 4 and lower row of cyst 5. A) benzbromarone m/z = 422.9061, B) overlay of benzbromarone m/z = 422.9061 (green) and PE(38:4) m/z = 766.5391 (blue), C) benzbromarone m/z = 422.9061 and hydroxybenzbromarone m/z = 438.9008 (red). Analysis was performed in negative ion mode with a step size of 50 µm, a scan time of 0.5s and a flow rate of 2 µL/min. The data was TIC normalized.

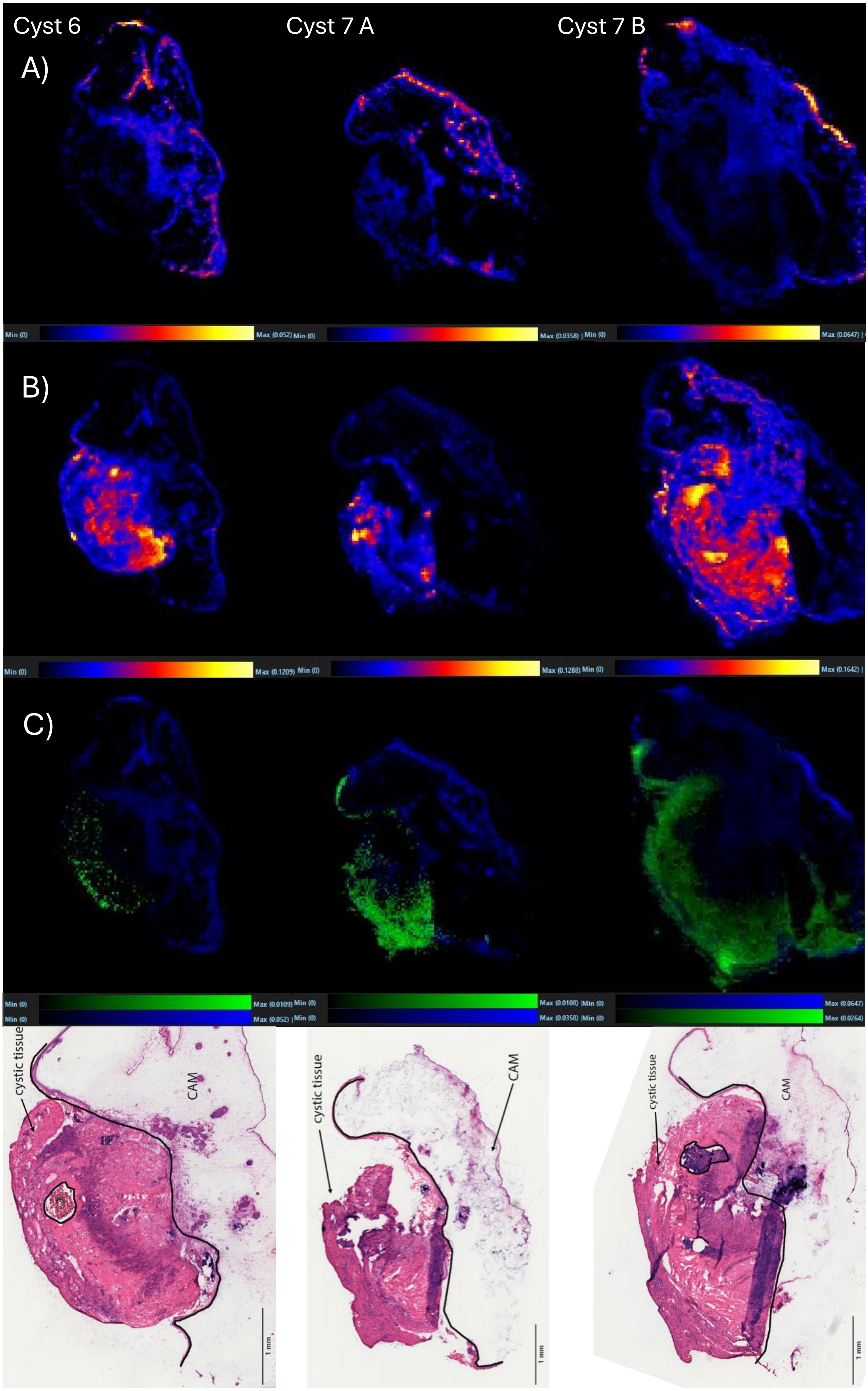
Cyst 6
Cyst 7 A
Cyst 7 B
A)
Min (0)
Max (0.052)
Min (0)
Max (0.0358)
Min (0)
Max (0.0647)
B)
Min (0)
Max (0.1209)
Min (0)
Max (0.1288)
Min (0)
Max (0.1642)
C)
Min (0)
Max (0.0109)
Min (0)
Max (0.052)
Min (0)
Max (0.0108)
Min (0)
Max (0.0358)
Min (0)
Max (0.0647)
Min (0)
Max (0.0264)
cystic tissue
CAM
1 mm
cystic tissue
CAM
1 mm
cystic tissue
CAM
1 mm

Supplementary Figure S3: HE and Ion images from two additional cyst samples (cyst 6 and Cyst 7 A/B) obtained from patient 3.

A) Ion image of PE 38:4 at m/z = 766.539 $[M-H]^-$
B) Ion image of Cer d34:1 at m/z = 572.481 $[M+Cl]^-$
C) Overlay of the ion images of Benzbromarone at m/z = 422.906 $[M-H]^-$ and PE 38:4 at m/z = 766.539 $[M-H]^-$
D) Corresponding HE stains. Note that for cyst 6 the consecutive section was annotated.

Mass spectral imaging analysis was performed in negative ion mode with a step size of 50 µm, a scan time of 0.5 s and a flow rate of 2 µL/min.

All data was TIC normalized.

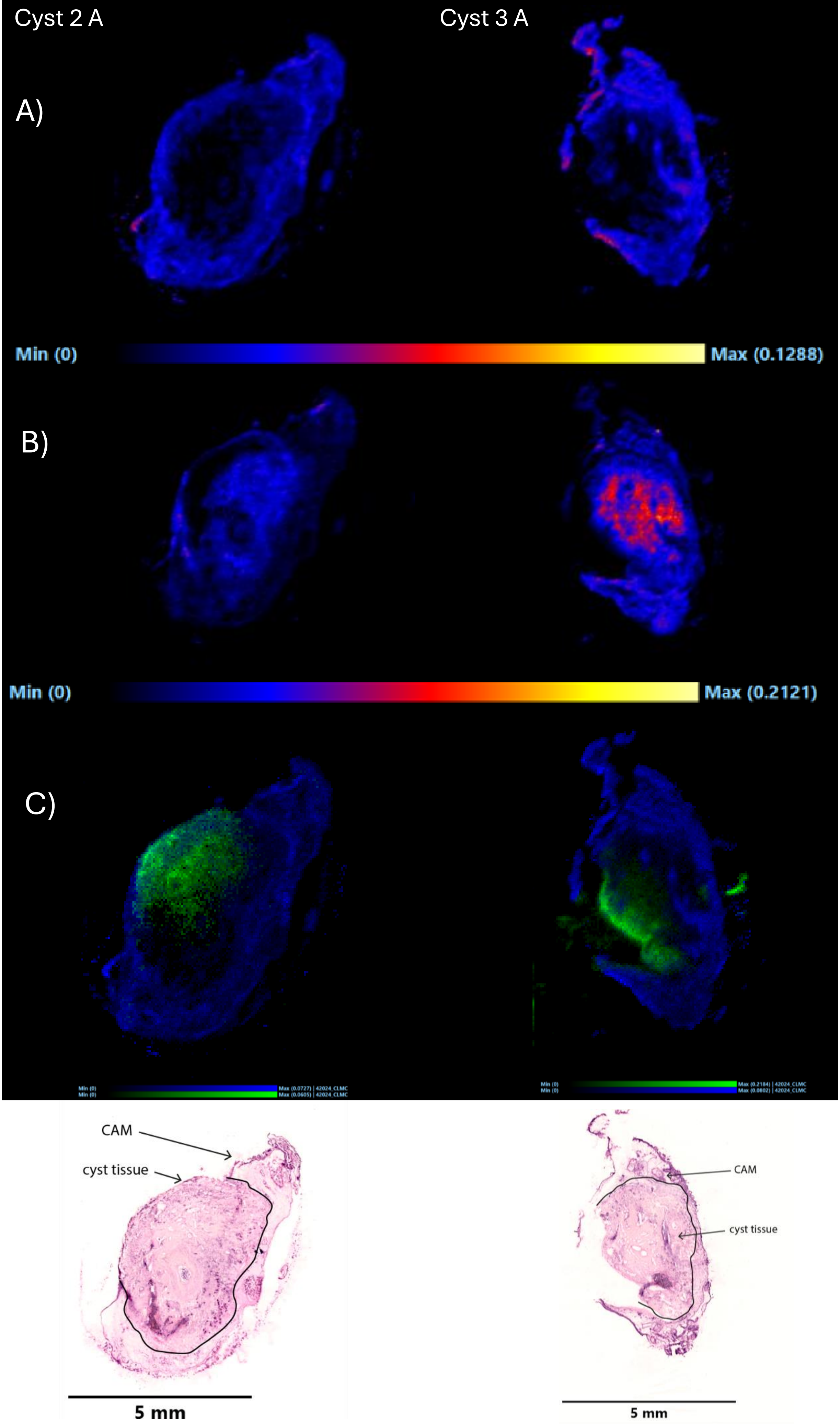

Cyst 2 A
Cyst 3 A
A)
Min (0)
Max (0.1288)
B)
Min (0)
Max (0.2121)
C)
Min (0)
Min (0)
Max (0.0727) | 42024_CLMC
Max (0.0605) | 42024_CLMC
Min (0)
Min (0)
Max (0.2184) | 42024_CLMC
Max (0.0802) | 42024_CLMC
CAM
cyst tissue
5 mm
CAM
cyst tissue
5 mm

Supplementary Figure S4: Ion images from cyst samples (Cyst 2 and cyst 10) obtained from patient 1.

A) Ion image of PE 38:4 at $m/z$ = 766.539 $[M-H]^-$
B) Ion image of Cer d34:1 at $m/z$ = 572.481 $[M+Cl]^-$
C) Overlay of the ion images of Benzbromarone at $m/z$ = 422.906 $[M-H]^-$ and PE 38:4 at $m/z$ = 766.539 $[M-H]^-$

Analysis was performed with a step size of 50 μm, a scan time of 0.5 s and a flow rate of 2 μL/min. All data was TIC normalized.

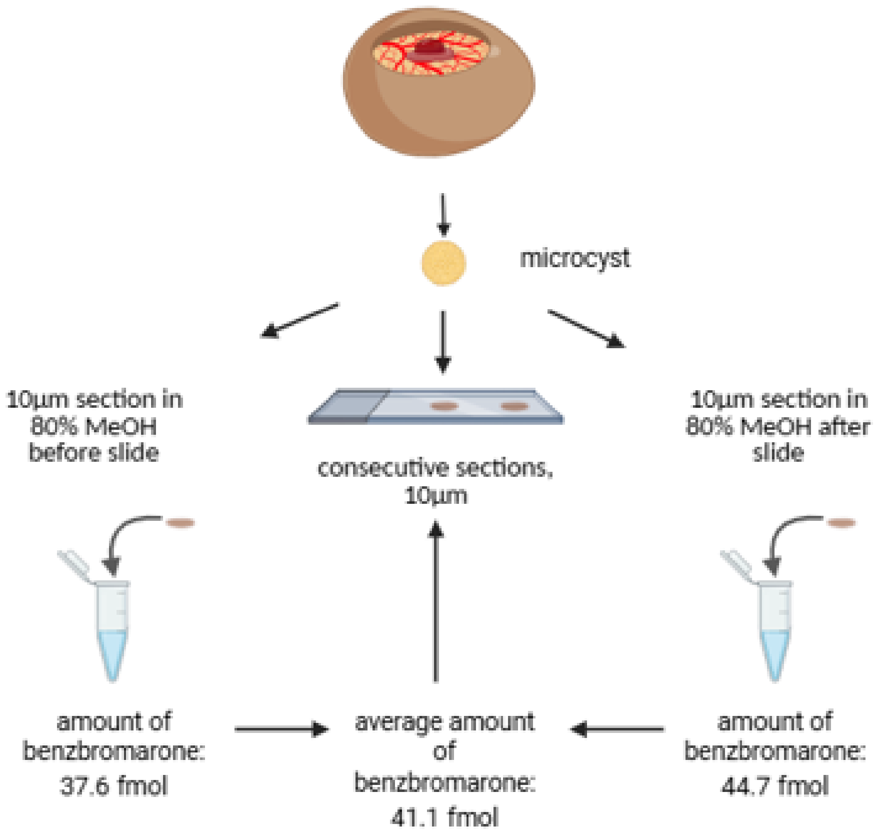


Supplementary Figure S5: Workflow employed for benzbromarone quantification.

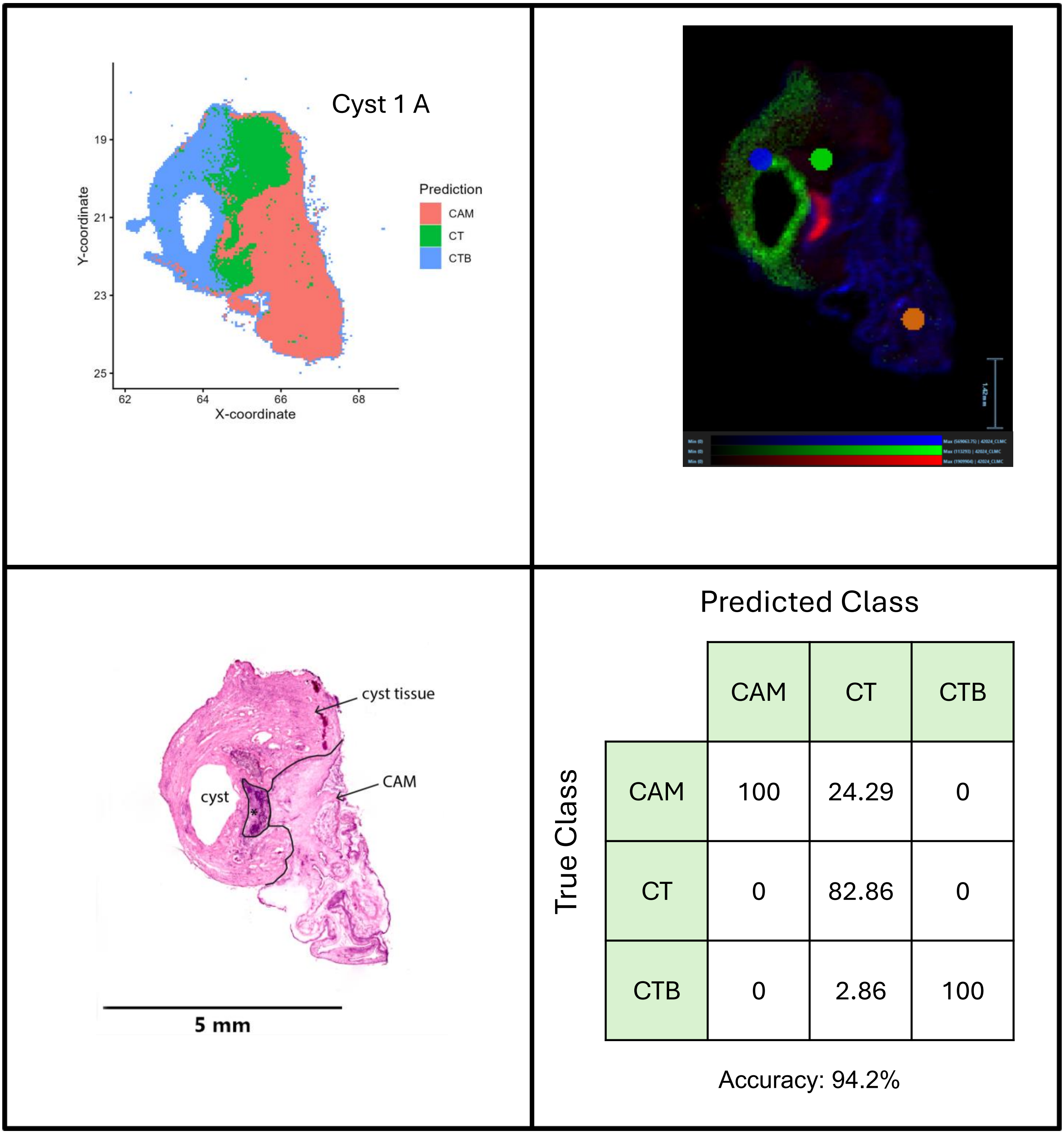


| | CAM | CT | CTB |
|---|---|---|---|
| CAM | 100 | 24.29 | 0 |
| CT | 0 | 82.86 | 0 |
| CTB | 0 | 2.86 | 100 |

Accuracy: 94.2%

Supplementary Figure S6: Results of the classification experiments for cysts 1A for patient 1. a) classification of CAM, CT, and CTB tissue areas using LASSO zero-sum signatures (1 – CAM, 2 – CT, 3 – CTB), b) an overlay of three signals (m/z = 422.960 (benzbromarone, green), m/z = 572.4824 (Cer d34:1, red), m/z = 766.5391 (PE38:4, blue). ROIs used for data analysis are shown in the overlayed ion image (orange: CAM, green: CT, blue: CTB), c) the HE stain of each cyst and d) the confusion matrix and the accuracy of the classification for each cyst.

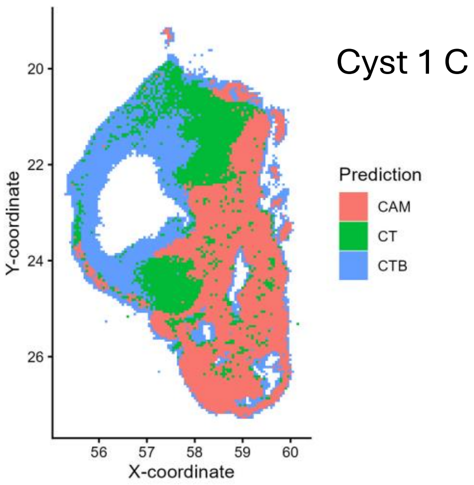


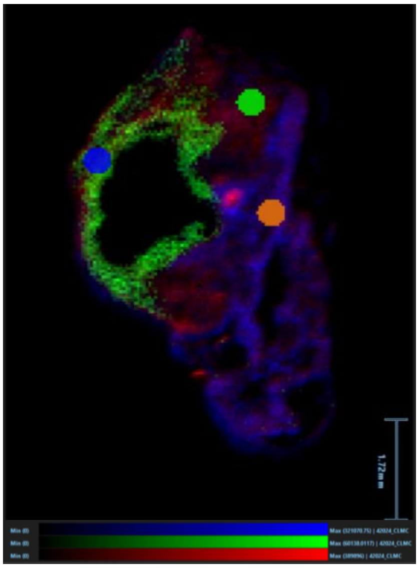

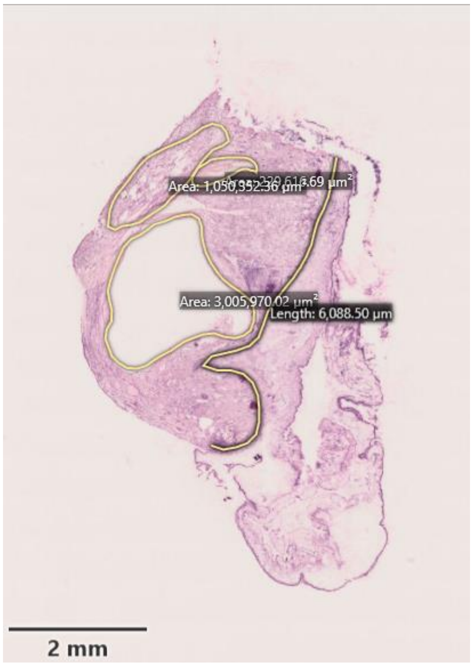


**Predicted Class** (columns) / **True Class** (rows)

| | CAM | CT | CTB |
|---|---|---|---|
| CAM | 98.57 | 0 | 0 |
| CT | 1.43 | 98.57 | 0 |
| CTB | 0 | 1.43 | 100 |

Accuracy: 99.04%

Supplementary Figure S7: Results of the classification experiments for cysts 1C for patient 1. a) classification of CAM, CT, and CTB tissue areas using LASSO zero-sum signatures (1 – CAM, 2 – CT, 3 – CTB, 4 ), b) an overlay of three signals (m/z = 422.960 (benzbromarone, green), m/z = 572.4824 (Cer d34:1, red), m/z = 766.5391 (PE38:4, blue). ROIs used for data analysis are shown in the overlayed ion image (orange: CAM, green: CT, blue: CTB), c) the HE stain of each cyst and d) the confusion matrix and the accuracy of the classification for each cyst.

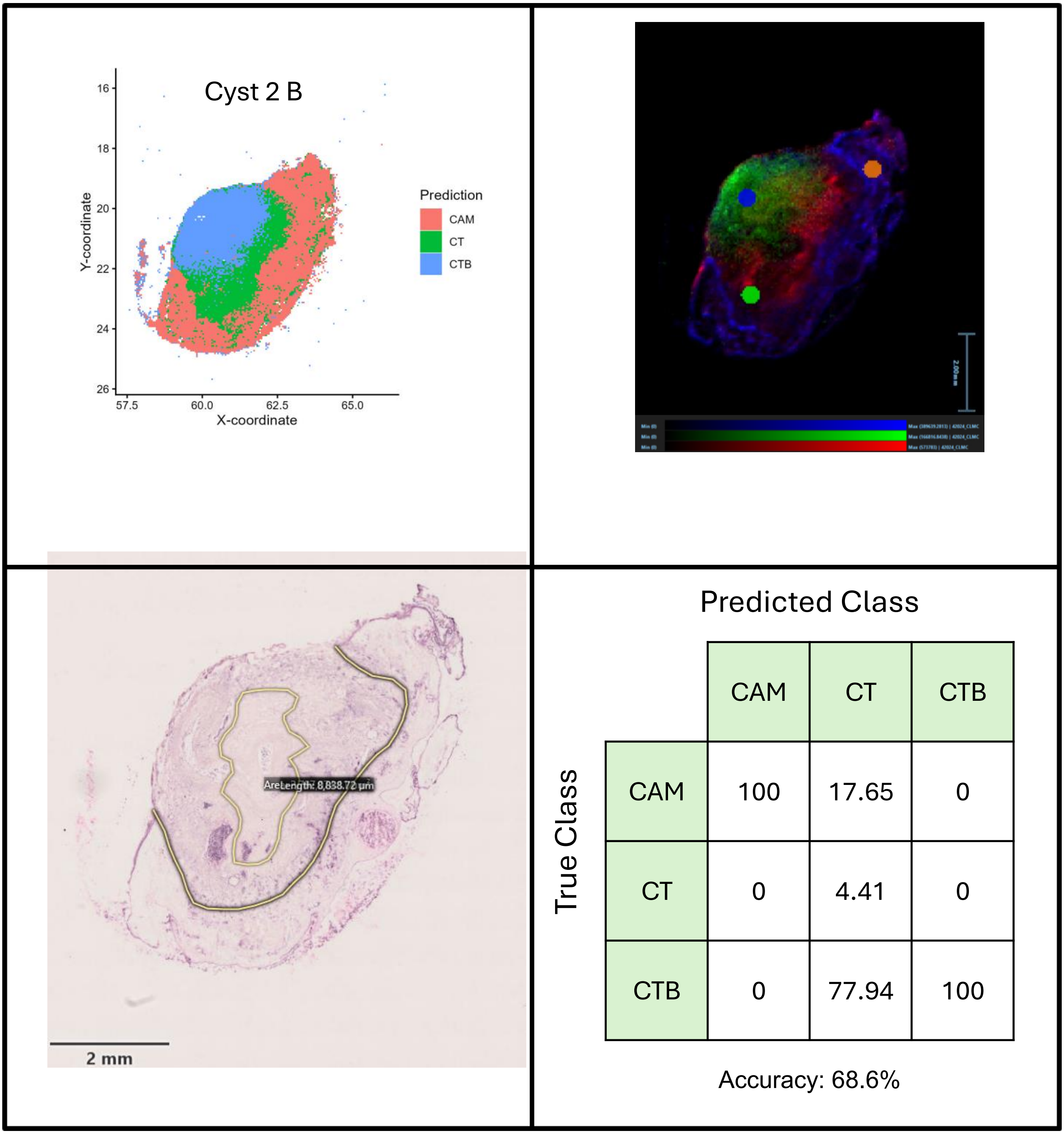


Supplementary Figure S8: Results of the classification experiments for cysts 2B for patient 1. a) classification of CAM, CT, and CTB tissue areas using LASSO zero-sum signatures (1 – CAM, 2 – CT, 3 – CTB, 4 ), b) an overlay of three signals (m/z = 422.960 (benzbromarone, green), m/z = 572.4824 (Cer d34:1, red), m/z = 766.5391 (PE38:4, blue). ROIs used for data analysis are shown in the overlayed ion image (orange: CAM, green: CT, blue: CTB), c) the HE stain of each cyst and d) the confusion matrix and the accuracy of the classification for each cyst.

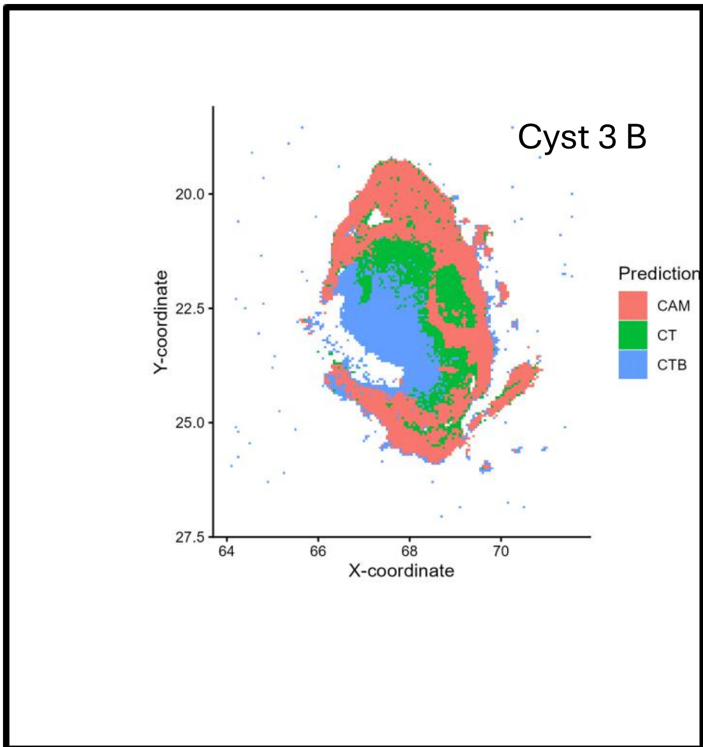


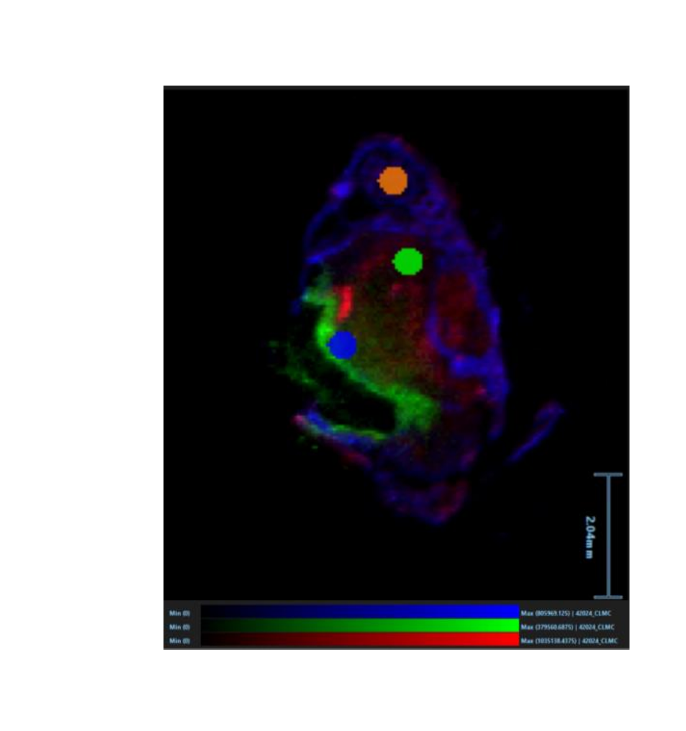


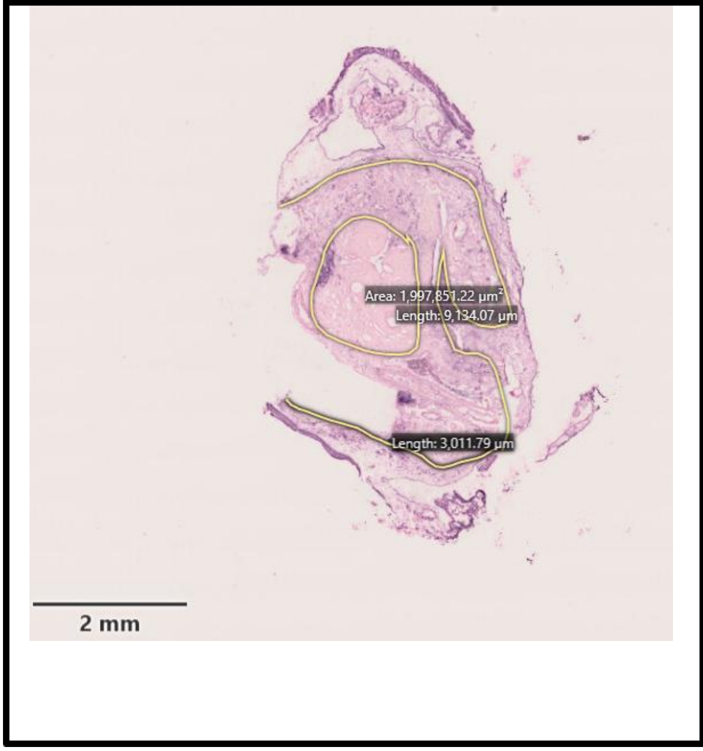


Predicted Class

| True Class | CAM | CT | CTB |
|---|---|---|---|
| CAM | 100 | 4.48 | 0 |
| CT | 0 | 95.52 | 0 |
| CTB | 0 | 0 | 100 |

Accuracy: 98.55%

Supplementary Figure S9: Results of the classification experiments for cysts 3B for patient 1. a) classification of CAM, CT, and CTB tissue areas using LASSO zero-sum signatures (1 – CAM, 2 – CT, 3 – CTB, 4 ), b) an overlay of three signals (m/z = 422.960 (benzbromarone, green), m/z = 572.4824 (Cer d34:1, red), m/z = 766.5391 (PE38:4, blue). ROIs used for data analysis are shown in the overlayed ion image (orange: CAM, green: CT, blue: CTB), c) the HE stain of each cyst and d) the confusion matrix and the accuracy of the classification for each cyst.

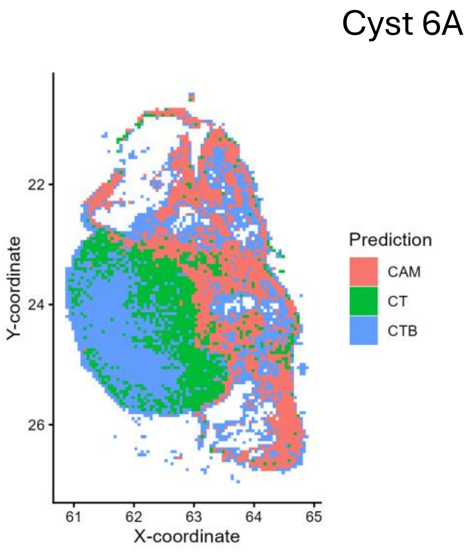


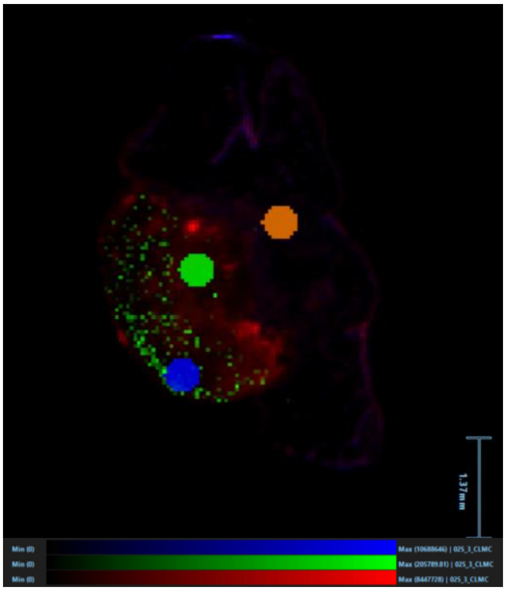

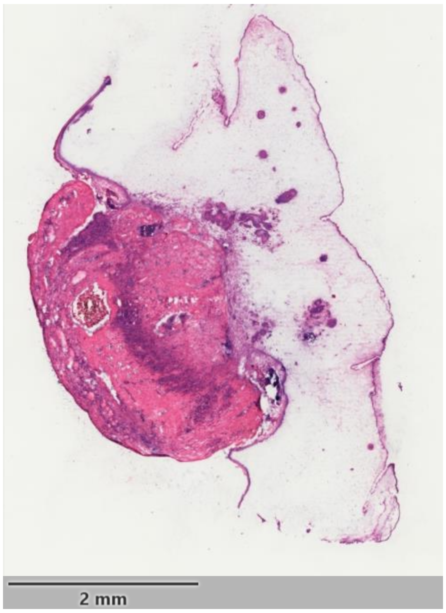


HE stain of consecutive slide

Predicted Class

| True Class | CAM | CT | CTB |
|---|---|---|---|
| CAM | 70 | 1.43 | 0 |
| CT | 30 | 42.86 | 0 |
| CTB | 0 | 55.71 | 100 |

Accuracy: 62.35%

Supplementary Figure S10: Results of the classification experiments for cysts 6A for patient 3. a) classification of CAM, CT, and CTB tissue areas using LASSO zero-sum signatures (1 – CAM, 2 – CT, 3 – CTB, 4 ), b) an overlay of three signals (m/z = 422.960 (benzbromarone, green), m/z = 572.4824 (Cer d34:1, red), m/z = 766.5391 (PE38:4, blue). ROIs used for data analysis are shown in the overlayed ion image (orange: CAM, green: CT, blue: CTB), c) the HE stain of each cyst and d) the confusion matrix and the accuracy of the classification for each cyst.

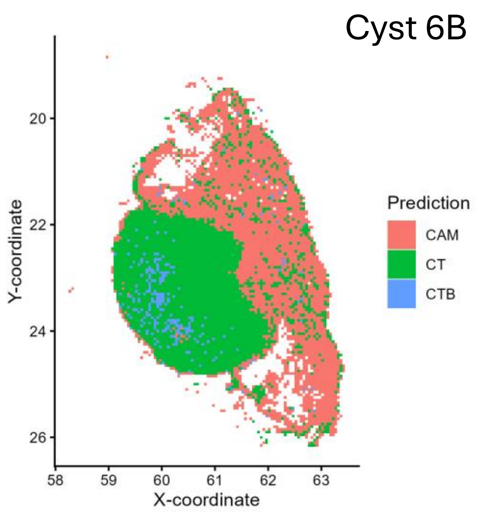


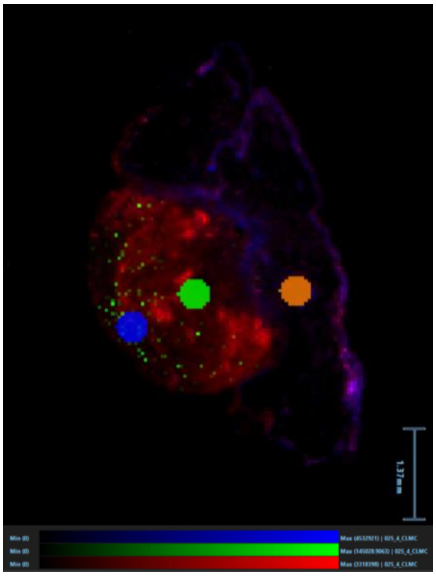

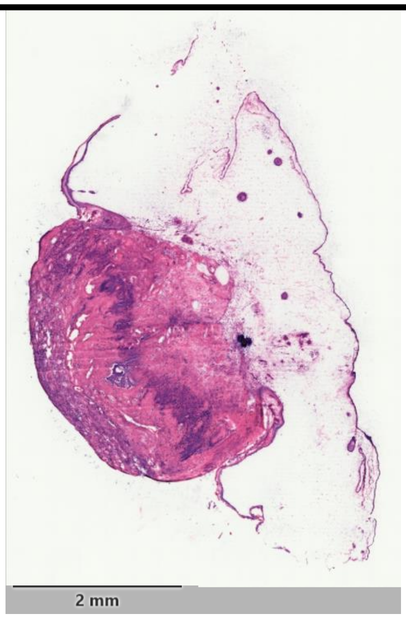


| | | Predicted Class | | |
|---|---|---|---|---|
| | | CAM | CT | CTB |
| True Class | CAM | 97.14 | 0 | 0 |
| | CT | 2.86 | 81.43 | 0 |
| | CTB | 0 | 18.57 | 100 |

Accuracy: 90.51%

Supplementary Figure S11: Results of the classification experiments for cysts 6B for patient 3. a) classification of CAM, CT, and CTB tissue areas using LASSO zero-sum signatures (1 – CAM, 2 – CT, 3 – CTB, 4 ), b) an overlay of three signals (m/z = 422.960 (benzbromarone, green), m/z = 572.4824 (Cer d34:1, red), m/z = 766.5391 (PE38:4, blue). ROIs used for data analysis are shown in the overlayed ion image (orange: CAM, green: CT, blue: CTB), c) the HE stain of each cyst and d) the confusion matrix and the accuracy of the classification for each cyst.

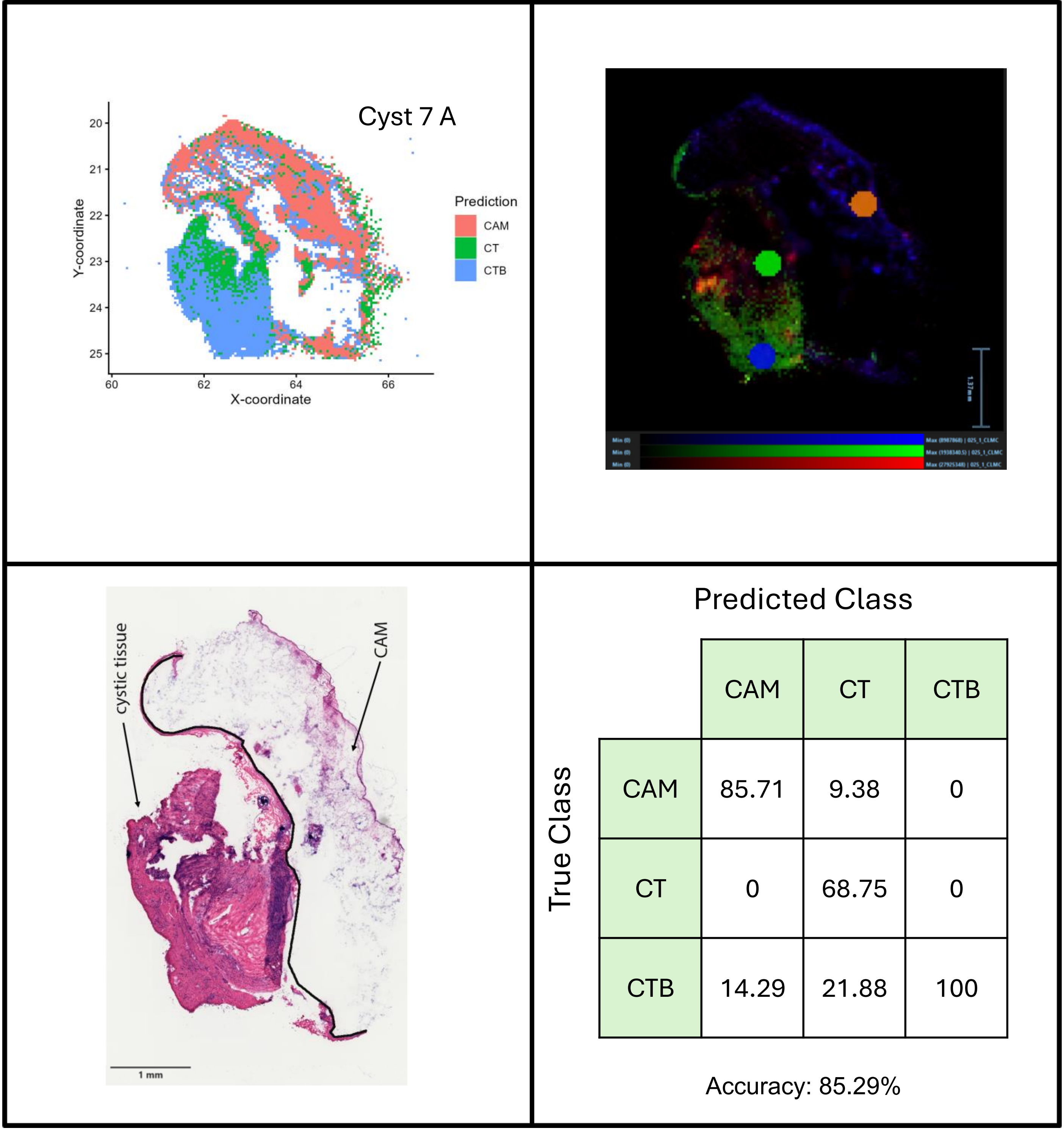


| | CAM | CT | CTB |
|---|---|---|---|
| CAM | 85.71 | 9.38 | 0 |
| CT | 0 | 68.75 | 0 |
| CTB | 14.29 | 21.88 | 100 |

Supplementary Figure S12: Results of the classification experiments for cysts 7A for patient 3. a) classification of CAM, CT, and CTB tissue areas using LASSO zero-sum signatures (1 – CAM, 2 – CT, 3 – CTB, 4 ), b) an overlay of three signals (m/z = 422.960 (benzbromarone, green), m/z = 572.4824 (Cer d34:1, red), m/z = 766.5391 (PE38:4, blue). ROIs used for data analysis are shown in the overlayed ion image (orange: CAM, green: CT, blue: CTB), c) the HE stain of each cyst and d) the confusion matrix and the accuracy of the classification for each cyst.

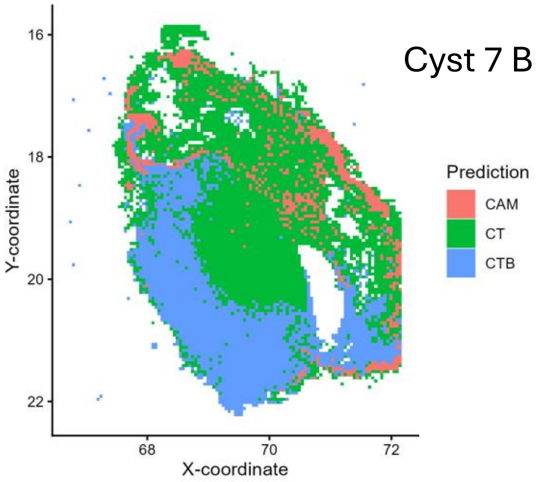


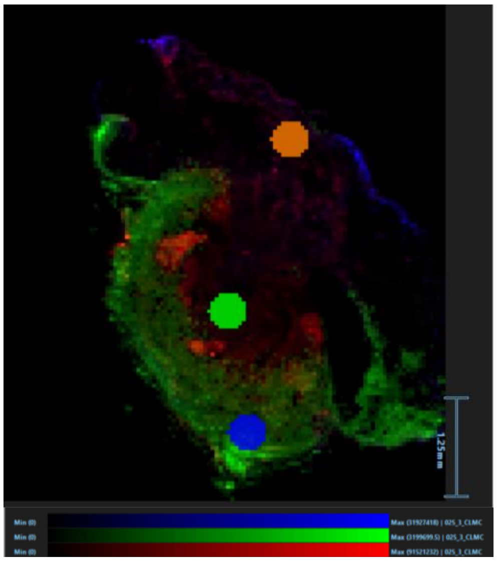

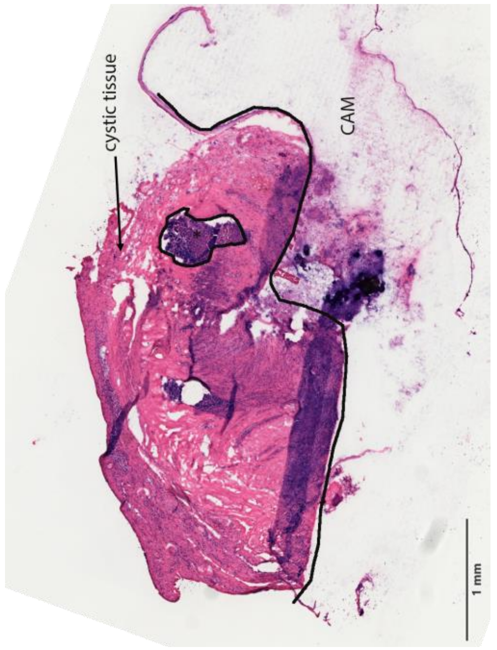


| True Class \ Predicted Class | CAM | CT | CTB |
|---|---|---|---|
| CAM | 28.57 | 0 | 0 |
| CT | 71.43 | 97.1 | 0 |
| CTB | 0 | 2.9 | 100 |

Accuracy: 75.12%

Supplementary Figure S13: Results of the classification experiments for cysts 7B for patient 3. 1. a) classification of CAM, CT, and CTB tissue areas using LASSO zero-sum signatures (1 – CAM, 2 – CT, 3 – CTB, 4 ), b) an overlay of three signals (m/z = 422.960 (benzbromarone, green), m/z = 572.4824 (Cer d34:1, red), m/z = 766.5391 (PE38:4, blue). ROIs used for data analysis are shown in the overlayed ion image (orange: CAM, green: CT, blue: CTB), c) the HE stain of each cyst and d) the confusion matrix and the accuracy of the classification for each cyst.

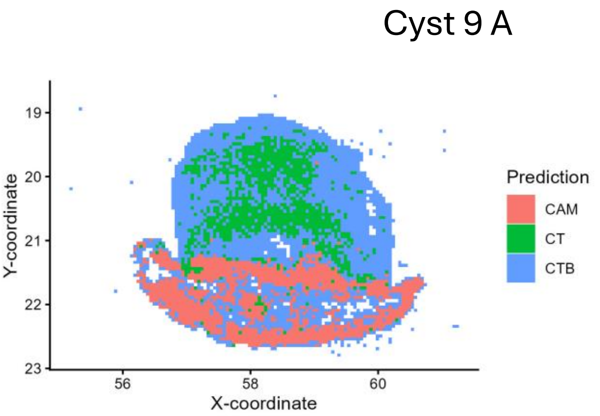


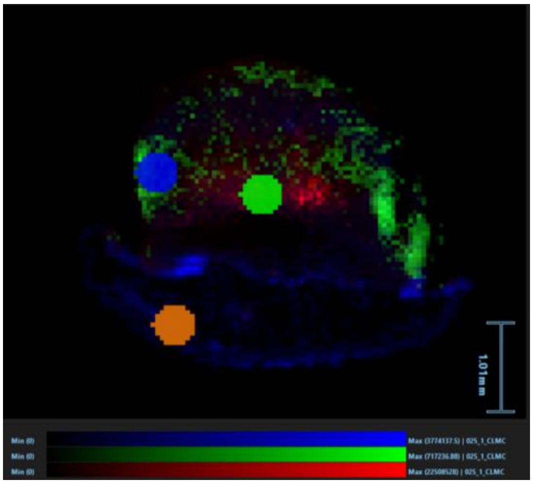


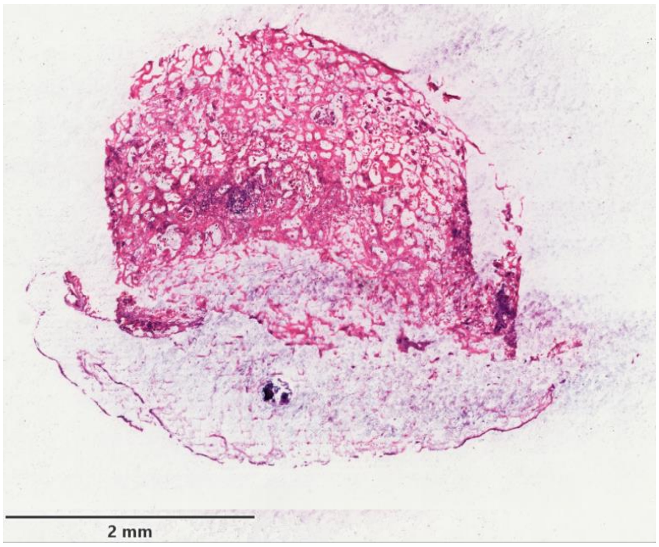


Predicted Class

| True Class | CAM | CT | CTB |
|---|---|---|---|
| CAM | 87.14 | 0 | 0 |
| CT | 0 | 43.08 | 0 |
| CTB | 12.86 | 56.92 | 100 |

Accuracy: 76.77%

Supplementary Figure S14: Results of the classification experiments for cysts 9A for patient 4. a) classification of CAM, CT, and CTB tissue areas using LASSO zero-sum signatures (1 – CAM, 2 – CT, 3 – CTB, 4 ), b) an overlay of three signals (m/z = 422.960 (benzbromarone, green), m/z = 572.4824 (Cer d34:1, red), m/z = 766.5391 (PE38:4, blue). ROIs used for data analysis are shown in the overlayed ion image (orange: CAM, green: CT, blue: CTB), c) the HE stain of each cyst and d) the confusion matrix and the accuracy of the classification for each cyst.

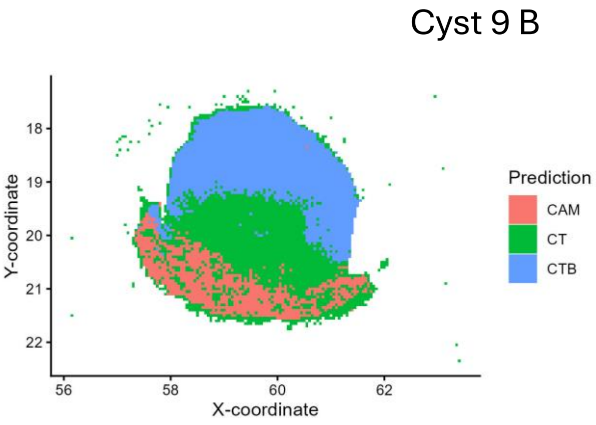


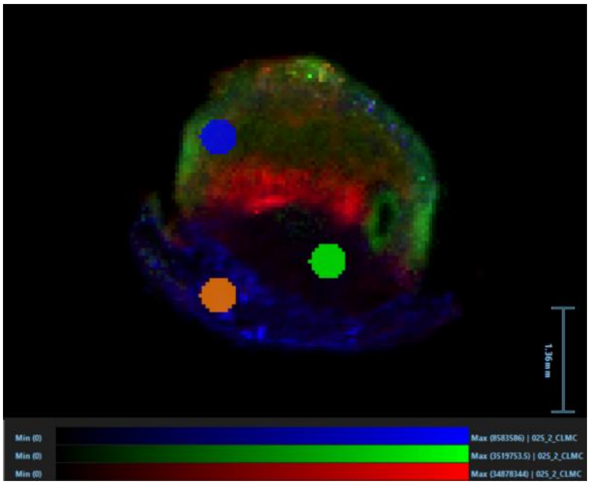

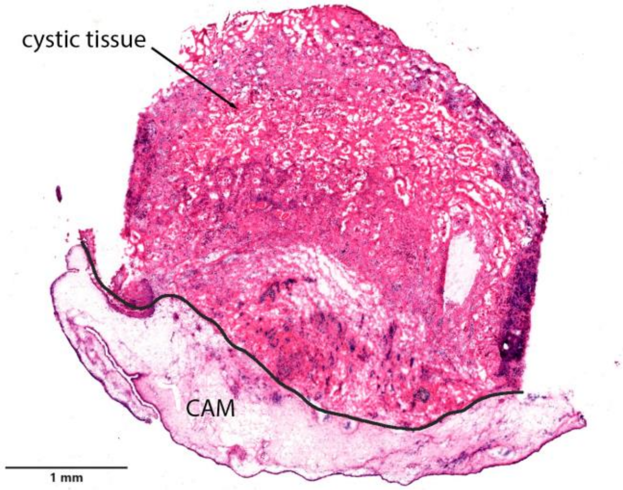


Predicted Class / True Class

| | CAM | CT | CTB |
|---|---|---|---|
| CAM | 100 | 24.29 | 0 |
| CT | 0 | 60 | 0 |
| CTB | 0 | 15.71 | 100 |

Accuracy: 86.67%

Supplementary Figure S15: Results of the classification experiments for cysts 9B for patient 4. a) classification of CAM, CT, and CTB tissue areas using LASSO zero-sum signatures (1 – CAM, 2 – CT, 3 – CTB, 4 ), b) an overlay of three signals (m/z = 422.960 (benzbromarone, green), m/z = 572.4824 (Cer d34:1, red), m/z = 766.5391 (PE38:4, blue). ROIs used for data analysis are shown in the overlayed ion image (orange: CAM, green: CT, blue: CTB), c) the HE stain of each cyst and d) the confusion matrix and the accuracy of the classification for each cyst.

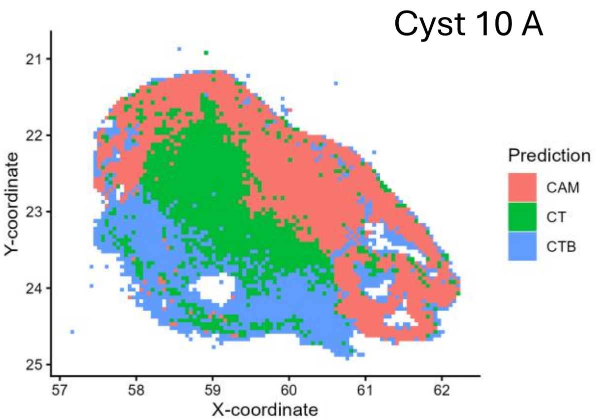


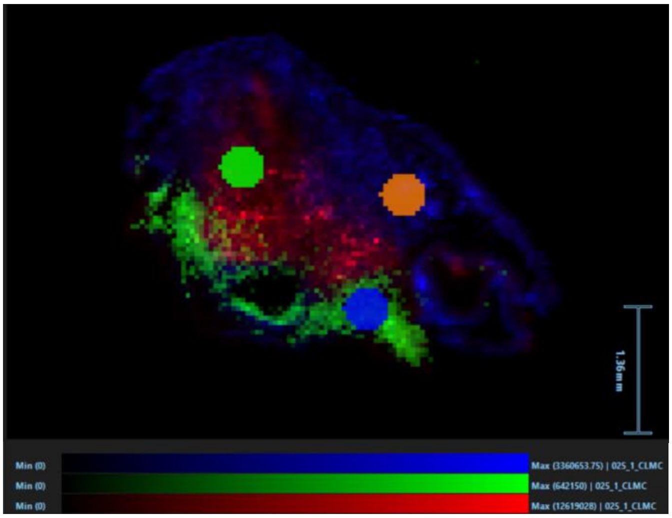

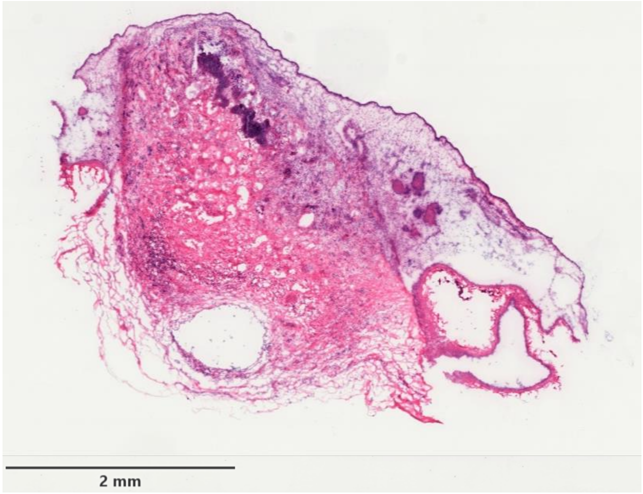


Predicted Class

| True Class | CAM | CT | CTB |
|---|---|---|---|
| CAM | 100 | 0 | 0 |
| CT | 0 | 82.86 | 0 |
| CTB | 0 | 17.14 | 100 |

Accuracy: 94.23%

Supplementary Figure S16: Results of the classification experiments for cysts 10A for patient 4. a) classification of CAM, CT, and CTB tissue areas using LASSO zero-sum signatures (1 – CAM, 2 – CT, 3 – CTB, 4 ), b) an overlay of three signals (m/z = 422.960 (benzbromarone, green), m/z = 572.4824 (Cer d34:1, red), m/z = 766.5391 (PE38:4, blue). ROIs used for data analysis are shown in the overlayed ion image (orange: CAM, green: CT, blue: CTB), c) the HE stain of each cyst and d) the confusion matrix and the accuracy of the classification for each cyst.

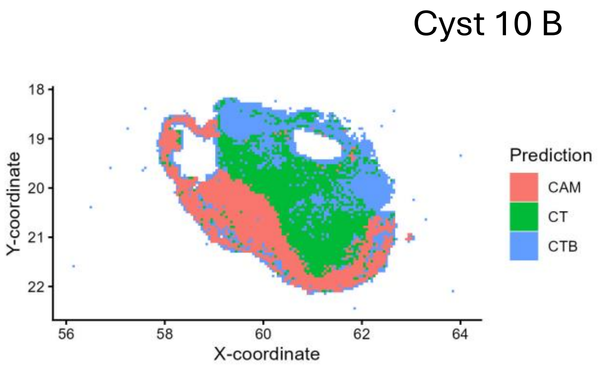


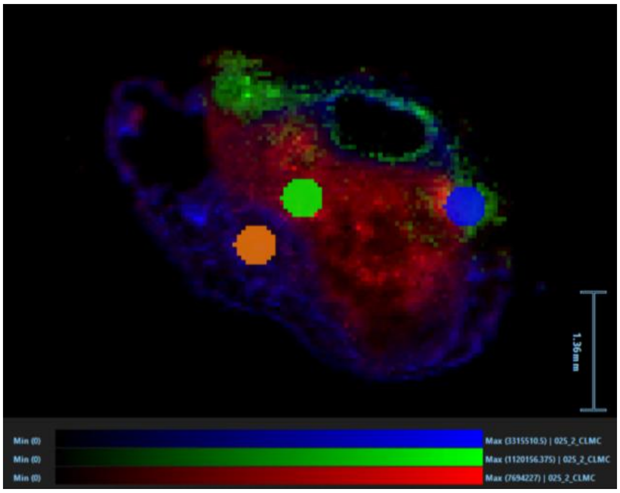


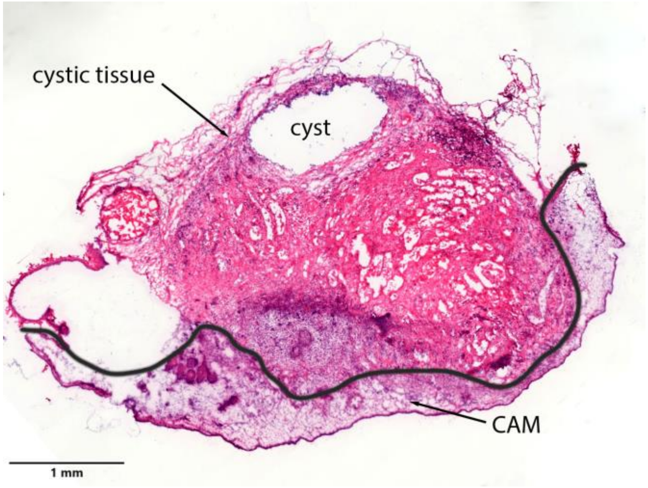


Predicted Class

True Class

| | CAM | CT | CTB |
|---|---|---|---|
| CAM | 97.14 | 0 | 0 |
| CT | 0 | 32.86 | 0 |
| CTB | 2.86 | 67.14 | 100 |

Accuracy: 76.56%

Supplementary Figure S17: Results of the classification experiments for cysts 10B for patient 4. a) classification of CAM, CT, and CTB tissue areas using LASSO zero-sum signatures (1 – CAM, 2 – CT, 3 – CTB, ), b) an overlay of three signals (m/z = 422.960 (benzbromarone, green), m/z = 572.4824 (Cer d34:1, red), m/z = 766.5391 (PE38:4, blue). ROIs used for data analysis are shown in the overlayed ion image (orange: CAM, green: CT, blue: CTB), c) the HE stain of each cyst and d) the confusion matrix and the accuracy of the classification for each cyst.

**Supplementary Table S1: Samples analyzed. 17 sections stemming from 10 cysts and 4 patients were measured in total.**

| Cyst sample | Section | dataset | calssification | quantification | patient |
|---|---|---|---|---|---|
| 1 | A | cyst 1A-1 | training | no quantification | 1 |
| | A | cyst 1A-2 | training | no quantification | |
| | A | cyst 1A-3 | testing | no quantification | |
| | B | cyst 1B | testing | no quantification | |
| | C | cyst 1C | testing | no quantification | |
| 2 | A | cyst 2A | testing | no quantification | |
| | B | cyst 2B | testing | no quantification | |
| 3 | A | cyst 3A | testing | no quantification | |
| | B | cyst 3B | testing | no quantification | |
| 4 | A | cyst 4 | not used for training/testing | no quantification | 2 |
| 5 | A | cyst 5 | not used for training/testing | no quantification | |
| 6 | A | cyst 6A | testing | quantification | 3 |
| | B | cyst 6B | testing | quantification | |
| 7 | A | cyst 7A | testing | no quantification | |
| | B | cyst 7B | testing | no quantification | |
| 8 | A | cyst 8A | testing | quantification | |
| | B | cyst 8B | testing | quantification | |
| 9 | A | cyst 9A | testing | no quantification | 4 |
| | B | cyst 9B | testing | no quantification | |
| 10 | A | cyst 10A | testing | no quantification | |
| | B | cyst 10B | testing | no quantification | |

**Supplementary Table S2: Non-zero coefficients of the multinomial zero-sum LASSO model.**

| Measured m/z | Coefficient CAM | Coefficient CT | Coefficient CTB | Calculated m/z | Formula neutral | Putative identification | \|Δ m\| | Ion |
|---|---|---|---|---|---|---|---|---|
| 215.0325 | | 0.2466624 | -0.2205525 | 215.0328 | C6H12O6 | Hexose, FA 6:0;O4 | 0.0003 | [M+Cl]- |
| 217.0296 | | 0.6153356 | -0.4751613 | 217.0298 | C6H12O7 | Hexose, FA 6:0;O5 (37Cl isotope) | 0.0002 | [M+Cl]- |
| 255.2328 | | 0.175457 | -0.0159614 | 255.233 | C16H32O2 | FA 16:0, FAL 16:0;O, SFE 16:0, WE 16:0 | 0.0002 | [M-H]- |
| 422.906 | | -0.7824002 | 1.5771192 | 422.906 | C17H12Br2O3 | Benzbromarone ($^{81}Br_1$ isotope) | 0 | [M-H]- |
| 528.2731 | | 0.0453798 | | | | | | |
| 570.4657 | | 0.0316292 | | 570.4658 | C34H65NO3 | Palmitoyl oxostearamide | 0.0001 | [M+Cl]- |
| 626.5364 | -0.3271271 | | | | | | | |
| 627.5397 | -0.0927056 | | | | | $^{13}C_1$ isotope of m/z 626.5364 | | |
| 640.552 | | 0.1778364 | -0.2151595 | | | | | |
| 682.5909 | -0.0312585 | 0.0441507 | | 682.5911 | C42H81NO3 | Cer 42:2;O2 | 0.0002 | [M+Cl]- |
| 685.6099 | -0.0651254 | | | 685.6101 | C42H83NO3 | Cer 42:1;O2 ($^{13}C_1$ isotope) | 0.0002 | [M+Cl]- |
| 700.5285 | | 0.1454773 | | 700.5287 | C39H76NO7P | CerP 39:2;O3; LPC 31:2; LPC O-31:3;O; LPE 34:2; LPE O-34:3;O; PE O-34:2 | 0.0002 | [M-H]- |
| 716.5234 | | -0.0873972 | | 716.5235 | C39H76NO8P | PE 34:1 | 0.0001 | [M-H]- |
| 737.5365 | | 0.0220877 | | | | | | |
| 738.5078 | 0.2295656 | -0.4128148 | | 738.5079 | C41H74NO8P | CerP 41:5;O4; LPC 33:5;O; LPT O-34:5; PE 36:4; PE O-36:5;O | 0.0001 | [M-H]- |
| 742.5391 | 0.3301371 | | | 742.5392 | C41H78NO8P | CerP 41:3;O4; LPC 33:3;O; LPT O-34:3; PE 36:2; PE O-36:3;O | 0.0001 | [M-H]- |
| 744.5547 | | 0.0713732 | | | | | | |
| 767.5426 | | | -0.3321597 | 767.5426 | C43H78NO8P | CerP 43:5;O4; PE 38:4; PE O-38:5;O ($^{13}C_1$ isotope) | 0.0001 | [M-H]- |
| 768.5314 | | -0.020227 | | | | | | |
| 768.5457 | | -0.0373176 | | | | | | |
| 771.5181 | | -0.5031 | | 771.5182 | C42H77O10P | PG 36:3; PG O-36:4;O | 0 | [M-H]- |

| | | | | | | | | |
|---|---|---|---|---|---|---|---|---|
| 773.5336 | | | -0.0678043 | | | | | |
| 774.5372 | 0.4393609 | | | | | $^{13}C_1$ isotope of m/z 773.5335 | | |
| 774.5441 | 0.0447401 | | | 774.5443 | C45H78NO7P | PE O-40:7 | 0.0002 | [M-H]- |
| 778.5755 | | | -0.2503203 | | | | | |
| 779.5789 | 0.1615668 | | | | | $^{13}C_1$ isotope of m/z 778.5754 | | |
| 786.5289 | | -0.060101 | | 786.529 | C42H78NO10P | PS(36:2 ) | 0.0001 | [M-H]- |
| 788.5446 | -0.2805658 | 0.0950923 | | 788.5447 | C42H79NO10P | PS(36:1) | 0.0001 | [M-H]- |
| 789.548 | -0.4085881 | 0.3105042 | | | | $^{13}C_1$ isotope of m/z 788.5445 | | |
| 790.539 | | -0.0776284 | | 790.5392 | C45H78NO8P | PE 40:6; PE O-40:7;O | 0.0002 | [M-H]- |
